# Deletion of TNFR1 in astrocytes restores memory in aged Alzheimer’s disease mice

**DOI:** 10.64898/2026.04.02.716130

**Authors:** Toko Kikuchi, Ioannis Zalachoras, Julien Prados, Alexis Assens, Roberta De Ceglia, Manuel Mameli, Ludovic Telley, Andrea Volterra

## Abstract

Astrocytes participate in local inflammation and cognitive decline in Alzheimer’s disease (AD). Aberrant cytokine TNFα signaling via astrocyte type-1 receptor (aTNFR1) could causally link the two AD pathology aspects. To verify this hypothesis, we crossed transgenic AD mice with mice enabling astrocyte-specific conditional TNFR1 deletion (aTNFR1KO). Induction of aTNFR1KO at early AD stages, preserved memory and reduced β-amyloid load and astrogliosis in the aged mice. Induction of aTNFR1KO at late AD stages, in mice already memory-impaired, surprisingly produced rapid memory rescue, without affecting β-amyloid load and astrogliosis. Single nucleus-RNA-seq analysis of all hippocampal cell populations revealed that late-stage aTNFR1KO rapidly modifies gene expression mainly in neurons, primarily targeting synaptic pathways, causing combined glutamatergic downregulation and GABAergic up-regulation. Consistently, hippocampal EEGs showed a pro-inhibitory effect of aTNFR1KO, which thus restores memory by “rebalancing” hippocampal circuitry excitability. This pro-memory effect identifies a new mechanism and astrocyte target against cognitive decline in AD.

## Introduction

Increasing evidence indicates that astrocytes play key pathogenic roles in Alzheimer’s disease (AD). Notably, in a population of aged subjects with β-amyloid deposits but no cognitive symptoms, increased blood levels of the astrocyte cytoskeletal protein, glial fibrillary acidic protein (GFAP), identify those subjects that will take a trajectory towards cognitive decline and AD^1,2^. This observation has important clinical implications that make GFAP a promising early AD biomarker^3^.At the transcriptional level, appearance of abnormal astrocyte populations is another trait that distinguishes those subjects that deviate from normal aging towards AD and cognitive decline^2,4^. Increased GFAP levels signal a state of reactive astrogliosis that drives astrocytes, together with microglia, into noxious local inflammatory loops in cognitive regions^5,6^. Among the mediators released at these inflammatory loci, the cytokine tumour necrosis factor-α (TNFα) may be deleterious and contribute to AD pathology^7^. Under physiological conditions, glia-originating TNFα plays regulatory roles in the synaptic circuitry and governs homeostatic synaptic plasticity, for example by controlling insertion of AMPA and GABA_A_ receptor subunits at post-synaptic membranes in an activity-dependent manner^8–11^. We found that TNFα also controls glutamate release from astrocytes^12^, which in turn strengthens the release probability of hippocampal excitatory synapses^13^, and contributes to their long-term plasticity and to contextual memory^14^. Under inflammation, increased ambient TNFα levels can, however, alter the above processes and thereby produce relevant circuital dysfunctions^15–18^. Notably, we showed that increased TNFα causes excess glutamate release from astrocytes with neuro-damaging consequences^19^. In a mouse model of multiple sclerosis, enhanced levels of TNFα at local foci of leukocyte infiltration in the dorsal hippocampus, induce overactivation of astrocyte TNFα type-1 receptor (aTNFR1), resulting in persistently increased glutamatergic transmission and impaired memory^20^. Local inflammatory foci with reactive astrocytes and microglia are present around β-amyloid plaques in AD. Thus, we became interested in directly assessing whether altered aTNFR1 signaling plays a memory-damaging role in AD. Towards this goal, we developed a new quadruple transgenic mouse line obtained by crossing an AD line, 5xFAD, which recapitulates several human AD traits including memory deficits^21,22^ and transcriptional alterations^23^, with a line that we generated to enable conditional TNFR1 deletion selectively in astrocytes (aTNFR1KO) (Methods). The resulting combined line allowed us deleting aTNFR1 at different stages of the AD pathology and evaluate the impact on memory, notably at late stages, when the mice are already impaired, i.e. in the window of opportunity for therapeutic interventions.

## Results

### AD mice with inducible astrocyte-specific TNFR1 deletion

To verify if our planned aTNFR1 mouse studies were potentially relevant to human AD, we initially analysed five existing human AD transcriptomic databases and, in all of them, found that aTNFR1 expression was up-regulated, like in 5xFAD mice, confirming that alteration of this astrocyte receptor is a common trait of the AD pathology in AD mice and humans (**Table 1**). Next, to specifically address if dysfunctional aTNFR1 contributes to cognitive impairment in AD, we crossed female^24^ *5xFAD^tg/+^*mice (5xFAD) with sex-matched *GFAPCreERT2^tg/+^tdTomato^Isl/Isl^TNFR1^fl/fl^* mice, a new line in which the floxed TNFR1 gene can be conditionally deleted in astrocytes upon tamoxifen-induced (TAM) *Cre* recombination driven by the GFAP promoter, and in which the red reporter td-Tomato (tdTOM) is expressed in parallel in the recombined cells (**Fig. 1a,b**). The latter line was obtained by crossing *GFAPCreERT2^tg/+^tdTomato^Isl/Isl^* ^25^ with *TNFR1^fl/fl^* mice^26^. To study pathology progression in AD mice carrying aTNFR1 deletion with respect to AD mice without the deletion, we used three 5xFAD mouse line genotypes and different treatments: (1) *5xFAD^tg/+^GFAPCreERT2^tg/+^ tdTomato^Isl/Isl^TNFR1^fl/fl^* mice treated with TAM, i.e. the KO mice that, from now on, we will call for simplicity AD-aTNFR1KO mice (**Fig. 1a**); (2) *5xFAD^tg/+^ GFAPCreERT2^tg/+^tdTomato^Isl/Isl^ TNFR1^fl/fl^* treated with vehicle (Veh), i.e. the same AD genotype as (1) but in which we do not activate *Cre* recombination. We will call these mice AD Veh; and (3) TAM-treated *5xFAD^tg/+^tdTomato^Isl/Isl^ TNFR1^fl/fl^* mice, in which lack of the *GFAPCreERT2^tg/+^* construct prevents TAM-dependent *Cre* recombination, but allows control for unspecific TAM effects (**Fig. 1b**). We will call this group the AD TAM mice. In our experiments, we also used non-transgenic 5xFAD littermates (*5xFAD^+/+^*) carrying the *GFAPCreERT2^tg/+^tdTomato^Isl/Isl^ TNFR1^fl/fl^* construct. When treated with Veh, these mice represented our non-AD control group (called CTRL), whereas, when treated with TAM, causing aTNFR1 deletion, they were called CTRL-aTNFR1KO mice and used to verify AD-unrelated effects of the aTNFR1 deletion. A summary of the steps involved in the generation of the above mouse lines as well as their original and simplified nomenclature and the color-coding used for identifying them in figures is presented in **Fig. 1a**.

**Figure 1.**
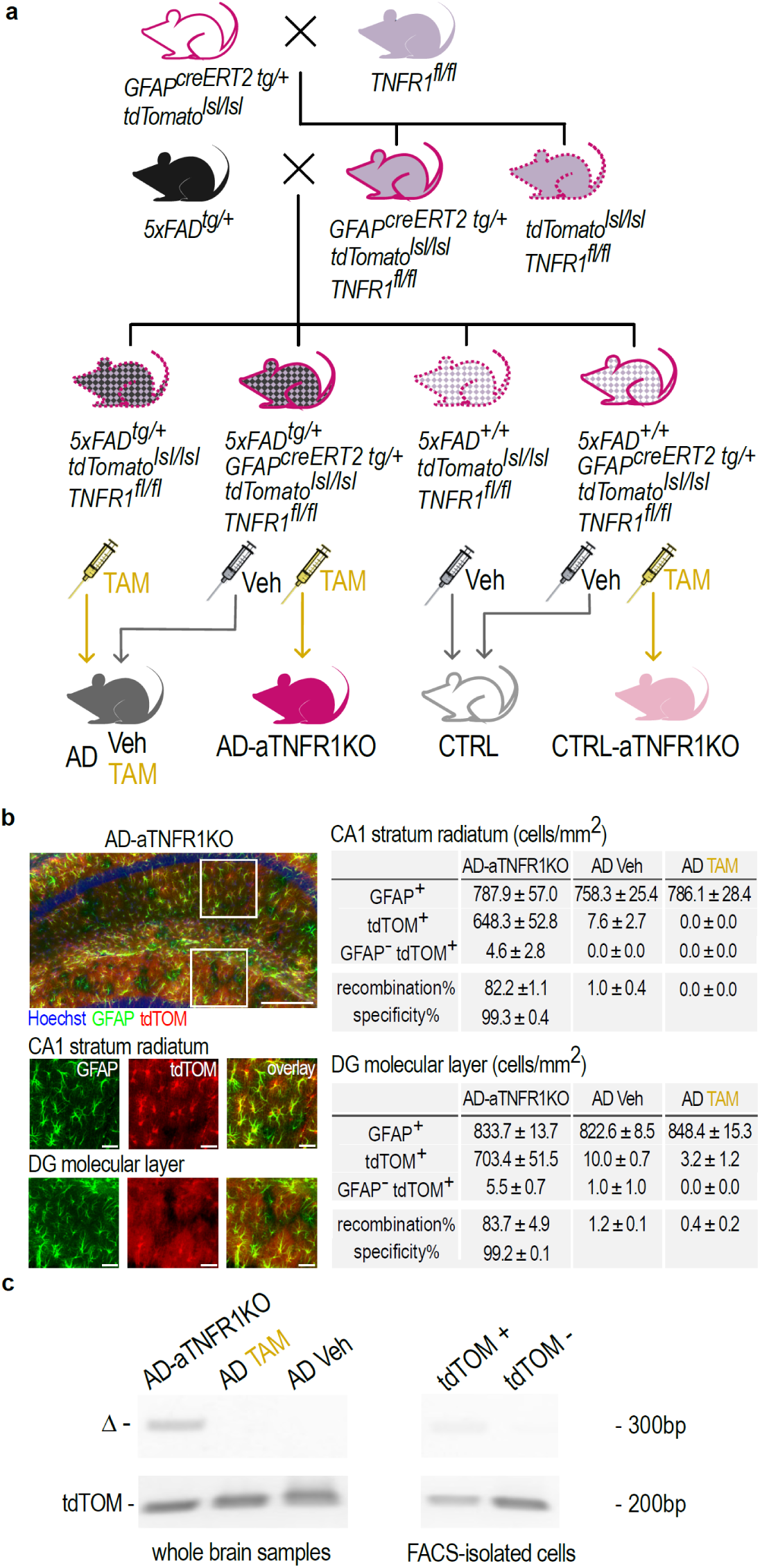
Generation and characterization of astrocyte-specific TNFR1KO mice with a 5xFAD background and related experimental lines. **a**. Breeding scheme for the generation of our experimental groups, including AD and non-AD (CTRL) mice, with or without astrocyte-specific TNFR1 deletion. Transgenic mice with two floxed *TNFR1* alleles (*TNFR1^fl/fl^*, mouse indigo) were crossed with heterozygous mice expressing tamoxifen (TAM)-inducible *Cre* recombinase under the astrocyte GFAP promoter and containing two *Cre*-inducible STOP-flox alleles of the red reporter *tdTomato* (tdTOM) (*GFAP^creERT2tg/+^tdTomato^lsl/lsl^,* white mouse with magenta contour) to produce *GFAP^creERT2tg/+^tdTomato^lsl/lsl^TNFR1^fl/fl^* (indigo mouse with magenta contour) and *tdTomato^lsl/lsl^TNFR1^fl/fl^* (indigo mouse with dotted magenta contour) offspring. The offspring were in turn crossed with *5xFAD^tg/+^* mice (black mouse) to produce *5xFAD^tg/+^GFAP^CreERT2tg/+^tdTomato^lsl/lsl^TNFR1^fl/fl^* mice (squared black-indigo mouse with magenta contour) and wild-type littermate *5xFAD^+/+^GFAP^CreERT2tg/+^tdTomato^Isl/lsl^TNFR1^fl/fl^* mice (squared white-indigo mouse with magenta contour), in which TNFR1 could be selectively knocked out in astrocytes following TAM treatment, producing the AD-aTNFR1KO (magenta mouse) and CTRL-TNFR1KO (pink mouse) experimental groups, respectively. In addition, crossing of the above offspring with *5xFAD^tg/+^* mice produced littermates lacking the *GFAP^CreERT2tg/+^* construct, i.e. *5xFAD^tg/+^tdTomato^lsl/lsl^TNFR1^fl/fl^* mice (squared black-indigo mouse with dotted magenta contour) and corresponding wild-types, i.e. *5xFAD^+/+^tdTomato^lsl/lsl^TNFR1^fl/fl^* mice (squared white-indigo mouse with dotted magenta contour): the former, treated with TAM, gave the AD TAM group (grey mouse). Instead, *5xFAD^tg/+^GFAP^CreERT2tg/+^tdTomato^lsl/lsl^TNFR1^fl/fl^* mice treated with Veh gave the AD Veh group (also shown as grey mouse). Together, they were used to investigate the 5xFAD-dependent effects, and, given the analogous results (**Extended data, Fig. 2**), they were merged and are collectively referred to as the AD group. In relation to the AD-TNFR1KO group, AD TAM mice were used to control for TAM-induced *Cre*-unrelated effects, and AD Veh for *Cre* leakage. Finally, *5xFAD^+/+^GFAP^CreERT2tg/+^tdTomato^lsl/lsl^TNFR1^fl/fl^* and *5xFAD^+/+^tdTomato^lsl/lsl^TNFR1^fl/fl^* mice treated with Veh were used as non-pathological controls, forming the CTRL group (white mouse with grey contour). **b,** *left*: representative images showing robust and selective *Cre*-dependent *tdTomato* (tdTOM, red) reporter expression in hippocampal astrocytes (GFAP-positive, green) of 9 months-old AD-aTNFR1KO mice treated with TAM (Fig. 2a**, *left***). Top: global image of the hippocampus (blue: Hoechst nuclear staining; white square insets are CA1 and DG regions shown below at higher magnification); middle: zoom-in image snapshots of astrocytes (green), tdTOM-positive cells (red) and their overlay in the CA1 stratum radiatum; bottom: same zoom-ins in the molecular layer of the dentate gyrus (DG). Scale bars = 200 µm for the global hippocampal image and 40 µm for zoomed images (white squares insets in the large images). *Right:* quantification of tdTOM-positive cells in the CA1 stratum radiatum and DG molecular layer of 9 month-old TAM-treated AD-aTNFR1KO mice (n=6 slices, 3 mice), as well as in AD Veh (n=4 slices, 2 mice) and AD TAM mice (n=4 slices, 2 mice). Data are presented as mean ± SEM. In AD-aTNFR1KO mice, *Cre*-dependent tdTOM expression was observed in >80% of GFAP-positive astrocytes with >99% specificity. In the absence of *Cre* recombinase expression (AD TAM mice) or TAM treatment (AD Veh mice) virtually no tdTOM expression was observed. **c.** Detection of *TNFR1* locus genetic deletion (Δ band) in whole brain homogenates of AD-aTNFR1KO, but not in homogenates of AD TAM and AD Veh mice, as well as in FACS-isolated tdTOM-positive cells from AD-aTNFR1KO, but not in tdTOM-negative cells. Genomic tdTOM bands were detected in the same samples to confirm the presence of genomic DNA (full gel scans are provided in the **Supplementary Data**).

**Table 1.** Astrocyte TNFR1 expression upregulation in AD patients and mouse models. Adjusted p-values are based on false-discovery rate. HC: hippocampus, PFC: prefrontal cortex, EC: entorhinal cortex.

| Study | Organism | Region | Astrocyte TNFR1:<br>% increase vs Control | Adjusted<br>p-value |
| --- | --- | --- | --- | --- |
| Mathys *et al*, 2024 * | Human | HC | 18.9 | 4.8E-22 |
| Sadick *et al*, 2022 † | Human | PFC | 26.5 | 1.3E-234 |
| Leng *et al*, 2021 ‡ | Human | EC | 29.3 | 1.0E-08 |
| Lau *et al*, 2020 § | Human | PFC | 10.1 | 6.0E-05 |
| Grubman *et al*, 2019 ¶ | Human | EC | 93.3 | 3.0E-14 |
| Habib *et al*, 2020 # | Mouse | HC | 26.1 | 1.0E-13 |

\* Comparison based on histopathological criteria (NIA-Reagn scale) – Reference: 58
† Comparison between AD patients and control subjects - Reference: 59
‡ Comparison between Braak stage 0 and 2 (early pathology) - Reference: 27
§ Comparison between AD patients and control subjects - Reference: 60
¶ Comparison between AD patients and control subjects - Reference: 61

To start, we quantified efficacy and specificity of TAM-induced *Cre* recombination in two different hippocampal regions, CA1 stratum radiatum and molecular layer of dentate gyrus in 9 months-old mice (**Fig. 1b**). While in AD (TAM or Veh) mice we saw almost no tdTOM-positive cells, confirming lack of unspecific recombination, in AD-aTNFR1KO mice, >80% of the GFAP-positive astrocytes were also tdTOM-positive, indicating high efficacy of recombination. To ascertain actual TNFR1 gene excision, we performed genomic PCR from both whole brain samples and FACS-isolated extracts from the same mice as above (**Fig. 1c**). In brain samples, we identified a band corresponding to the excised floxed TNFR1 gene sequence, selectively in AD-aTNFR1KO mice, and in these mice, we found the same band only in tdTOM-positive FACS-isolated cells. Overall, these data (as well as data later presented in **Extended data, Fig. 4f**) confirm astrocyte-selective TNFR1 deletion under our experimental conditions.

### AD pathology in our experimental transgenic lines

To study the impact of aTNFR1 deletion on pathology progression in AD mice, we performed studies in 9 month-old animals, i.e. at an age when our 5xFAD line shows plateau cognitive symptomatology^21,22^. We evaluated three main parameters: learning and memory performance, selecting as main behavioral test the contextual fear conditioning (CFC, **Extended data, Fig. 1a**, Methods), level of β-amyloid plaque deposition, and inflammatory reactive state of astrocytes (astrogliosis), the last two assessed in the hippocampus via immunostaining with an anti-pan amyloid-β and an anti-GFAP antibody, respectively (details in Methods). To start, we confirmed that native 5xFAD mice, i.e., the parent line with no other transgene insertion, at this age show evident AD symptomatology when compared to age-matched littermate controls, consisting in: (a) reduced memory performance in the CFC test, notably in recalling the learned association between aversive stimuli and context in which the stimuli were received **(Extended data, Fig. 1b,c**). For this effect, we excluded via additional open field (OF, Methods) behavioural assessing, causes different from cognitive deficit, such as alterations in pain sensitivity, exploratory activity, motor function, and anxiety state of the mice (**Extended data, Fig. 1d-g**); (b) diffuse presence of β-amyloid deposits throughout the hippocampus, absent in the controls (**Extended data, Fig. 1h**), and (c) significant increase in the GFAP-immunopositive area of the hippocampus due to enhanced GFAP expression, typical of reactive astrocytes (**Extended data, Fig. 1i**).

Next, we checked if the AD lines prepared for our experimental investigations, deriving from the above native 5xFAD line via insertion of the GFAP-driven *CreERT2-Lox* system and/or the *tdTomato^Isl/Isl^TNFR1^fl/fl^* sequences, and stimulated via vehicle or TAM treatment, maintained the same AD phenotype observed in the parental line (**Extended data, Fig. 1**). In particular, we compared AD Veh and AD TAM lines (**Fig. 1a**) to *5xFAD^tg/+^* mice in the three above parameters selected to define the AD phenotype in 9 months-old mice. We found no significant difference among the three groups in any of those parameters (**Extended data, Fig. 2a-i**) and concluded that the genetic modifications introduced in the 5xFAD line do not affect AD pathology progression, making the derived lines well suited for studying the impact of aTNFR1 deletion on the AD phenotype. Moreover, lack of any difference between AD Veh and AD TAM mice, led us to merge their data in a single group simply called AD and representing the pathology phenotype against which we compared the observations in mice with induced aTNFR1 deletion (AD-aTNFR1KO).

### Early-stage aTNFR1 deletion preserves memory performance and reduces β-amyloid plaques and astrogliosis

Up-regulated TNFR1 expression in astrocytes is an alteration already present at early AD stages, both in 5xFAD mice^4^, and in AD patients^27^. Such stages largely overlap with the pre-symptomatic phases of the pathology preceding tangible cognitive alterations. Given our temporal control on the induction of aTNFR1 deletion in AD mice, we decided to initially administer TAM injections and induce the deletion in 2.5 months-old mice (early-stage aTNFR1KO), i.e., in mice displaying initial astrogliosis and β-amyloid deposits in the hippocampus but no cognitive alteration^21^. With this approach, we could study the impact of the KO on the course of the pathology, assessing the outcome when the mice reached 9 months of age (**Fig. 2a** *left*).

**Figure 2.**
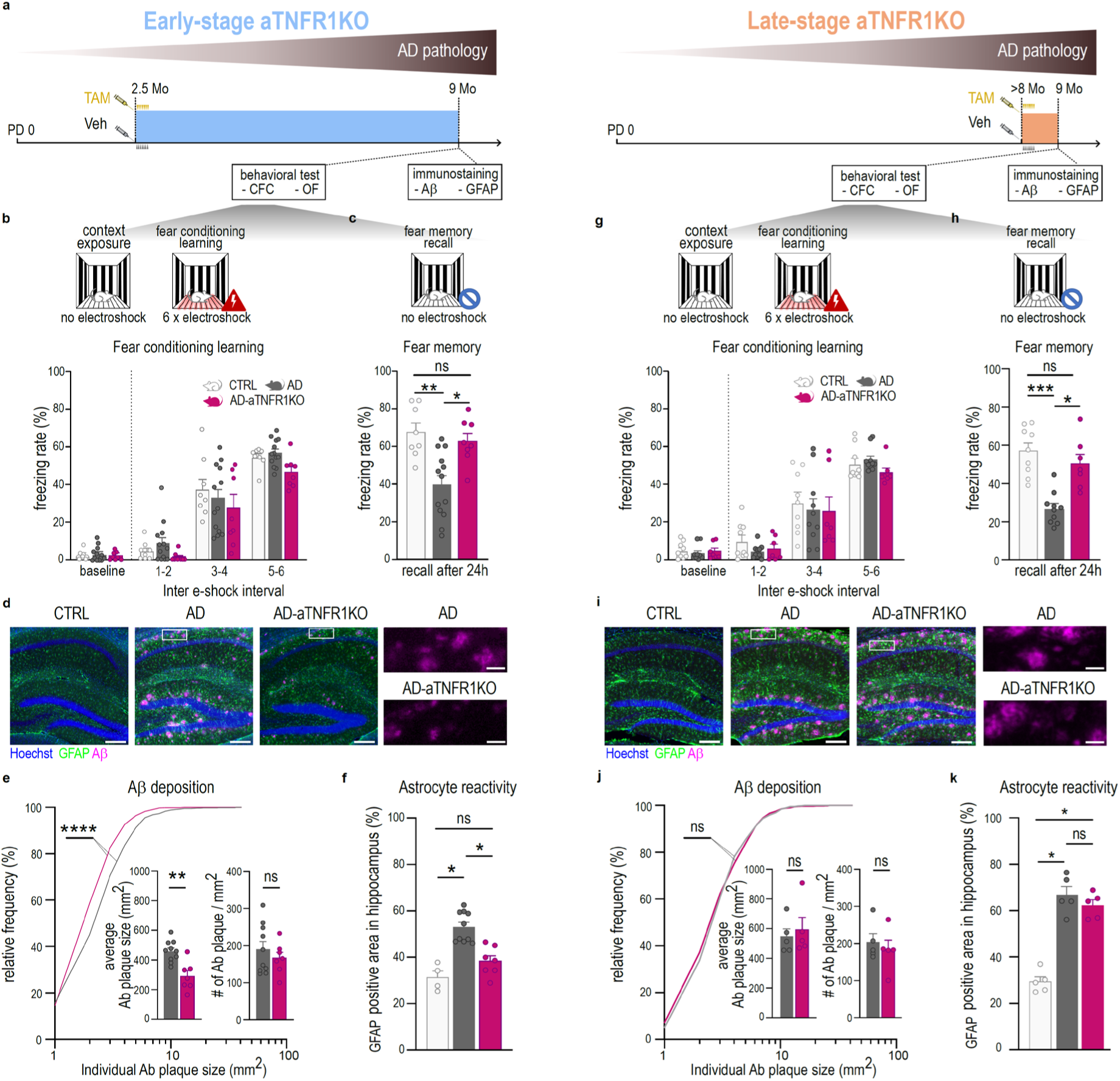
Astrocyte TNFR1 deletion at early or late AD pathology stages alleviates memory impairment, but has different effects on β-amyloid deposition and astrogliosis. **a.** Schematic representation of the experimental protocols: astrocytic TNFR1 deletion was induced at two different timepoints in independent AD mice cohorts (AD-aTNFR1KO). In the protocol on the *left*, deletion was induced via TAM injections at 2.5 months of age, before the emergence of memory deficits (early-stage aTNFR1KO, azure): while in the protocol on the *right,* deletion was induced at 8-8.5 months of age, when memory deficits have already appeared (late-stage aTNFR1KO, orange). In both protocols, two groups of AD mice (AD TAM and AD Veh) and the CTRL group were treated with the appropriate reagent (Veh or TAM, see Fig. 1a) and used to quantify AD pathology progression in comparison with the AD-aTNFR1KO group. In both protocols, all mouse groups were behaviourally tested at the age of ∼9 months in the open field (OF) and the contextual fear conditioning (CFC) tests. Brain tissue from the same mice was then collected post-mortem for β-amyloid (Aβ) and GFAP immunostaining. **b-f**: data from early-stage AD-aTNFR1KO protocol: **b.** Performance of the CFC test (Methods) in CTRL (white, N=9), AD (grey: AD TAM + AD Veh, see, **Extended data Fig. 2**, N=13) and AD-aTNFR1KO (magenta, N=8) mice. All groups showed similar performance during context exposure-baseline and fear conditioning (two-way ANOVA, group effect, p = 0.312; time x group interaction, p = 0.750). **c**. Fear memory was evaluated 24h after conditioning. AD mice showed lower freezing rates compared to CTRL indicating impaired fear memory, whereas AD-TNFR1KO mice showed higher freezing rates than AD mice indicating an amelioration of the AD-related memory deficits (Kruskal-Wallis test: H=11.60; p = 0.003, Dunn’s multiple comparisons test: AD vs AD-aTNFR1KO: p = 0.031; CTRL vs AD: p = 0.0068; CTRL vs AD-aTNFR1KO: p = 0.999). **d.** Representative images of the hippocampi of CTRL, AD and AD-aTNFR1KO mice showing β-amyloid (purple) GFAP (green) and cell nuclei (Hoechst, blue) immunoreactivity. Scale bars = 200 µm for large images, 40 µm for zoomed-in images (white rectangle insets in the large images). **e**, *left*: Quantitative comparison of Aβ plaque size frequency between AD and AD-aTNFR1KO mice. In AD-aTNFR1KO mice (N=7), there was a shift towards higher frequency of smaller Aβ plaques compared to AD mice (N=10) (Kolmogorov-Smirnov, D = 0.585; p < 0.0001). *Right:* bar plots show that the average size of Aβ deposits in AD-aTNFR1KO mice was significantly lower than in AD mice (Mann-Whitney two-tailed test, U = 4; p = 0.001), whereas the total number of the deposits was not different (Mann-Whitney two-tailed test, U = 31, p = 0.74). **f**. Area occupied by GFAP immunoreactivity in the hippocampus (Methods): the area in AD mice (N=10) was higher than in CTRL mice (N=4), indicating astrogliosis, but this was significantly reduced in AD-aTNFR1KO mice (N=7) and not different from CTRL mice (Kruskal-Wallis test, H = 15.56, p < 0.0001; CTRL vs AD: p = 0.01; AD, vs AD-aTNFR1KO: p = 0.012; CTRL vs AD-aTNFR1KO: p = 0.886). **g-k**: data from late-stage AD-aTNFR1KO protocol: **g.** Performance of the CFC test in CTRL (white, N=8), AD (grey, AD TAM + AD Veh, N=10) and AD-aTNFR1KO (magenta, N=7) mice. All groups showed similar performance during context exposure-baseline and CFC (Two-way ANOVA, group effect, p = 0.544; time x group interaction, p = 0.846). **h**. Fear memory evaluation 24 h after the conditioning: the AD group showed lower performance compared to CTRL, while AD-aTNFR1KO mice had significantly higher performance compared to AD mice, and not different from CTRL, indicating recovery from AD-related memory deficits despite the KO induction occurs in already memory impaired mice (Kruskal-Wallis test: H=16.09; p = 0.0003, Dunn’s multiple comparisons test: AD vs AD-aTNFR1KO: p = 0.0134; CTRL vs AD: p = 0.0004; CTRL vs AD-aTNFR1KO: p = 0.999). **i**. Representative images of the hippocampi of CTRL, AD and AD-aTNFR1KO mice showing β-amyloid (purple) GFAP (green) and cell nuclei (Hoechst, blue) immunoreactivity in the late-stage experiment. Scale bars as in **d**. **j**. Representation as in **e**. However, in the late-stage AD-aTNFR1KO experiment, Aβ deposits in AD-aTNFR1KO mice (N=5) are not significantly different from deposits in AD mice (N=5) in all parameters measured (relative frequency of the different sizes of the deposits: Kolmogorov-Smirnov, D = 0.244, p = 0.174; number of Aβ deposits: Mann-Whitney two-tailed test, U = 12, p = 0.99; average size of the Aβ deposits: Mann-Whitney two-tailed test, U = 9, p = 0.548). **k**. Quantification of GFAP immunoreactivity as in **f.** However, in the late-stage experiment, GFAP immunoreactivity was not significantly different between AD and AD-aTNFR1KO mice, and higher in both of these groups compared to CTRL mice (N=5 per group), (Kruskal-Wallis test, H = 9.500, p = 0.002: CTRL vs AD: p = 0.014; CTRL vs AD-aTNFR1KO: p = 0.04; AD vs AD-aTNFR1KO: p = 0.999).

To start, we evaluated whether early-stage aTNFR1KO influenced the development of the deficient memory phenotype present in the 9-month-old AD mice (**Extended data, Figs. 1 and 2**). For this, we compared the performance of AD-aTNFR1KO mice to that of AD mice as well as of their age-matched non-AD littermates (CTRL, **Fig. 2b,c**). Preliminarily, we assessed whether the three mouse groups showed any difference in exploratory or locomotor activity, and in their anxiety state that could act as confounders on the learning and memory performance. We did not observe any group differences (**Extended data, Fig. 3a-d**). During fear conditioning, all three mouse groups learned the association between context and aversive stimuli proficiently, exhibiting similar levels of conditioned fear at the end of the session **(Fig. 2b**). Moreover, all the groups moved the same distance during electroshocks (**Extended data, Fig. 3a**), excluding differences in pain sensitivity that could affect the CFC performance. However, when we measured contextual memory expression 24 h later, only AD mice exhibited a significant deficit with respect to the fear levels acquired during conditioning. In contrast, fear levels in AD-aTNFR1KO mice resembled those in control mice, indicating that induction of aTNFR1 deletion in young adult AD mice prevented pathological loss of memory when the mice aged (**Fig. 2c**). We then checked by immunohistochemistry in the mice subjected to behavioural testing, their levels of β-amyloid deposition (**Fig. 2d,e**) and reactive astrogliosis (**Fig. 2f**) in the hippocampus: we found that AD-TNFR1KO mice presented significantly reduced β-amyloid load compared to AD mice, notably showing plaques of smaller size. Likewise, the hippocampal area occupied by GFAP-positive signal, much larger in aged AD mice compared to controls, was reduced in AD-TNFR1KO mice to levels like controls. These results suggest that altered aTNFR1 function plays a primary role in the progression of the AD pathology because its deletion at early AD stages results in global attenuation, or slowing, of the pathology and, notably, prevents memory impairment.

### Late-stage aTNFR1 deletion rapidly improves memory without affecting β-amyloid plaques and astrogliosis

The above beneficial effect prompted us to test whether deletion of astrocyte TNFR1 could have a positive influence even when induced at late stages in the pathology, when 5xFAD mice manifest the plateau of their cognitive alterations^21,22,24^. For this, we induced aTNFR1KO in ∼8 months-old mice and evaluated learning and memory performance when the mice reached 9 months of age (late-stage aTNFR1KO, **Fig. 2a** *right*). Like in the previous tests with the early-stage protocol, all three mouse groups, CTRL, AD and AD-aTNFR1KO, subjected to the late-stage protocol performed well in learning the associative rule and becoming fear conditioned (**Fig. 2g**). During the contextual memory recall session 24 h later, only AD mice, however, showed the expected memory deficit. In contrast, AD-aTNFR1KO mice, once again, performed similarly to controls (**Fig. 2h**) and did not show any difference in potential confounding parameters relative to CTRL and AD mice (**Extended data, Fig. 3e-h**). Therefore, remarkably, late-stage aTNFR1KO leads to rescue of memory performance in already impaired mice. This recovery is even more impactful considering how short the period was between the receptor deletion and the behavioural testing, i.e. ∼3 weeks. Moreover, contrary to the observations in early-stage AD-aTNFR1KO, at immunohistochemical analysis, the hippocampi of late-stage AD-aTNFR1KO mice did not show any difference in AD neuropathology with respect to the corresponding 9 months-old AD mice without the deletion (**Extended data, Figs. 1 and 2**). Indeed, AD-aTNFR1KO mice were indistinguishable from AD mice in terms of overall deposition and size of β-amyloid plaques (**Fig. 2i,j**), and increased extent of GFAP-positive hippocampal area compared to CTRL mice (**Fig. 2k**). These data indicate that late-stage aTNFR1 deletion, like the early-stage knock-out, leads to improved memory performance. However, the beneficial effect of the late-stage deletion consists in a recovery of memory function, which is independent of actions on β-amyloid deposition and astrocyte inflammation, and thus involving a new, undefined, and rapidly acting mechanism.

### snRNA-seq transcriptional signature in our AD mice: correlation with human AD

To identify such mechanism, we decided to use an unbiased approach, performing wide-spectrum snRNA-seq transcriptional analysis of the biological processes modified by the late-stage deletion of astrocyte TNFR1 in the hippocampus. We thus compared the gene expression signature of all the neuronal and glial hippocampal cell populations of 9 months-old AD-aTNFR1KO mice to the signatures in age-matched AD mice and in their non-AD littermates, the latter with or without aTNFR1 deletion (**Fig. 3a**). Following quality control check (**Extended data, Fig. 4** and Methods), analysis of the transcriptional identities of the nuclei originating from our four experimental groups (Methods), revealed the presence of >20 transcriptionally defined clusters, which corresponded to the main hippocampal cell types. To specifically annotate each individual cluster, we trained a deep neural network model using a reference hippocampus-containing database (Methods).

**Figure 3.**
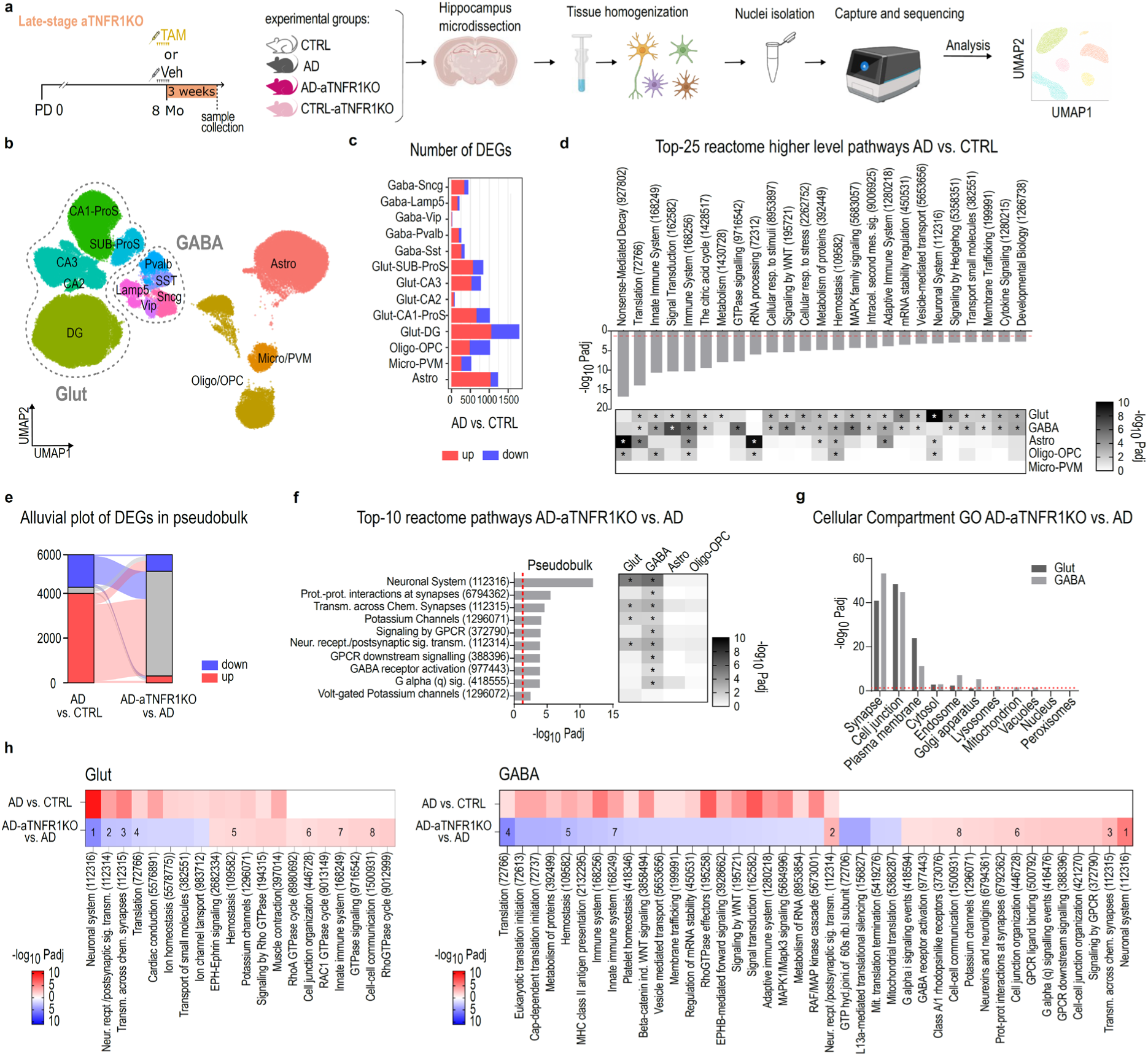
Rapid transcriptional changes induced by late-stage aTNFR1KO in AD mice indicate prominent effect on hippocampal glutamatergic and GABAergic neurons. **a**. Schematic representation of the experimental protocol. Mouse groups (CTRL, AD, AD-aTNFR1KO, and CTRL-aTNFR1KO) were treated with Veh or TAM as appropriate (Fig. 1a) at around 8 months of age, when AD mice are already memory-impaired, and aTNFR1KO is induced at late pathology stage. The mice were sacrificed 3 weeks later, their hippocampi microdissected, snap frozen and stored at -80C until further processing. Hippocampal tissue was homogenized for nuclei isolation, and subsequently the transcriptome of single nuclei was sequenced and analyzed. N=2-3 mice per group, split in two technical replicates. **b**. UMAP representation of the main identified hippocampal cell types based on snRNA-seq data. GABAergic and glutamatergic neuron subpopulations are grouped together with dashed lines. **c.** DEG between AD and CTRL mice and their distribution in the different cell types. Blue indicates down-regulation, red up-regulation. **d**. Top 25 differentially regulated biological pathways between AD and CTRL mice based on pseudobulk RNA expression and Reactome pathway enrichment analysis (Methods). Numbers in parentheses indicate unique Reactome pathway identifiers (they were added to enable certain identification of the pathways given that some names were abbreviated for space reasons). Histograms provide level of statistical significance in -log_10_Padj scale, with dotted red line signaling the significance threshold (1.3). The grey-scale heatmap below the histograms (also in log_10_Padj scale) shows the effect in each cell family and asterisks denote statistical significance (Methods). Only 1^st^ and 2^nd^ level Reactome pathways are shown in order to better capture the whole repertoire of AD-induced transcriptional changes. **e**. Alluvial plot showing the fate of hippocampal pseudobulk AD vs CTRL DEGs in the AD-aTNFR1KO vs AD comparison. Gene up-regulated (red), down-regulated (blue) or not modified (grey) in AD vs CTRL follow several patterns becoming up-regulated (red, a minority), down-regulated (blue, several) or being not modified (grey, the large majority) in AD-aTNFR1KO vs AD. **f**. Top10 differentially regulated biological pathways between AD-aTNFR1KO and AD mice in pseudobulk comparison with associated grey-scale heatmap indicating cell family-specific effects. Representation as in **d**, analysis as in **d**, with some difference (Methods). **G**. Cellular compartments with which top-ranked genes in glutamatergic (Glut) and GABAergic (GABA) neurons in the AD-aTNFR1KO vs AD comparison are associated (see Methods for analysis). “Cell junction”, “synapse” and “plasma membrane” are top 3 in both cell types, while the effect in all the other main cell compartments is much lower. **h.** Cell family-specific effects of AD-aTNFR1KO vs AD in glutamatergic (Glut) and GABAergic (GABA) neurons according to Reactome pathway enrichment analysis. All significantly regulated pathways for AD-aTNFR1KO vs AD are shown, or top 25 in each direction (up or down) if more than 25. Representation is as in **d**, except that the coloured heat-map scale identifies up-regulation (red) or down-regulation (blue). Moreover, the effects of AD vs CTRL are shown for the same pathways, allowing for direct comparison (e.g. if AD-aTNFR1KO vs AD affects a pathway also affected by AD vs CTRL or not – the latter represented in white in AD vs CTRL – and in which direction the AD-aTNFR1KO effect is, the same or opposite of the AD effect). The numbers in the heatmaps indicate pathways that were regulated by AD-aTNFR1KO vs AD in both Glut and GABA neurons, highlighting opposite regulation in some synapse-related pathways.

Cells annotated with an underrepresented cell type, with profiles suggesting contamination from cortical tissue, and low prediction confidence were removed (Methods). The final 13 predicted clusters encompassed different populations of glutamatergic and GABAergic neurons, as well as several non-neuronal cell types, including clusters identified as “astrocytes”, “oligo-OPC”, i.e., containing both mature and precursor cells of the oligodendroglial lineage, and “micro-PVM”, i.e., microglia and peri-vascular macrophages” (**Fig. 3b** and **Extended data, Fig. 4d,f,g**). These 13 clusters were present in all snRNA-seq samples obtained from the four experimental groups and showed limited differences in the respective proportions, thereby enabling cross-group comparisons (**Extended data, Fig. 4d,g**). Within the uniform manifold approximation and projection (UMAP) distribution, the astrocyte marker GFAP was enriched in the predicted “astrocyte” cluster (**Extended data, Fig. 4f**), and absent in all the neuronal ones, confirming correct prediction of our annotations as well as astrocyte-selectivity of late-stage TNFR1KO.

Based on this initial validation of our experimental samples, we went on to analyze transcriptional group differences, starting from those between AD mice and their age-matched controls (CTRL), representing the AD signature of our mouse line. Through differential gene expression analysis (Methods; **Fig. 3c** and **Supplementary Table 1**) we identified ∼8500 differentially expressed genes (DEGs) in AD mice with respect to CTRL mice, 60% of which were up-regulated and 40% down-regulated. DEGs were distributed throughout the hippocampal cell populations, but most abundantly (at least 1000 DEGs) in astrocytes, Oligo-OPC, and in Glutamatergic sub-populations of the CA1 (Glut-CA1-ProS) and dentate gyrus (Glut-DG) regions. To define the cellular processes and cell types most affected in AD mice, we conducted DEGs’ pathway enrichment analysis using the Reactome database^28^. To improve statistical power and correct imbalances in cell numbers between clusters, we aggregated for each sample our 13 cell clusters in a single transcriptomic profile (pseudobulk; see Ref.29 and Methods) and summarized statistics in specific grouped families of cell clusters (Glutamatergic neurons, GABAergic neurons, astrocytes, Oligo-OPC, Microglia-PVM). The top 25 cellular pathways significantly modified in AD mice (**Fig. 3d**), included, at the highest rankings, “control of gene expression”, “RNA translation”, “immune function”, “signal transduction” and “cell metabolism”. Pathway alterations were observed in all neuronal and non-neuronal cell populations and displayed cell family-specific patterns, except for microglia-PVM (**Fig. 3d** and **Supplementary Table 2**). To validate our observations and assess their relevance to other AD mouse models and, notably, to human AD, we then performed correlation analysis comparing our hippocampal DEGs dataset with analogous datasets available from previous studies in 5xFAD mice^4,30^, in other AD mouse lines (APP/PS1)^31^, as well as in AD patients^32,33^ (**Table 2**). Importantly, despite the methodological and biological differences across different studies, including the brain region of origin of the human datasets, our AD-related DEGs dataset showed significant correlation with all the others, including those from AD patients. Moreover, one of the human datasets^32^ contained information about DEGs in the different hippocampal cell populations, and this enabled us to confirm that the AD-related gene expression changes seen in each of our cell families in 5xFAD mice correlated with the changes in the corresponding human cell population. The only exception were the data in Microglia-PVM cells, which lacked significant correlation. For this reason, we decided to exclude this cell population from subsequent analyses. Overall, based on correlation analysis, the AD transcriptional signature identified in our 5xFAD mice is consistent with those observed in other AD mouse lines and in human AD, suggesting that our transcriptional observations could be of broad relevance, thus setting the stage for the further, more detailed analyses aimed at identifying the transcriptional changes associated with memory restoration in AD mice upon aTNFR1 deletion.

**Table 2.** Correlation of Alzheimer’s induced changes in gene expression between previous studies and the present dataset. Expression differences between control and 5xFAD mice in our dataset were correlated with expression differences between AD patients and controls or transgenic mice and their respective controls in each study. Pearson’s correlation coefficient (R) and p-values are provided. Green values denote significant correlations, whereas orange ones denote non-significant correlations. SEA-AD: Seattle Alzheimer’s Disease Brain Cell Atlas. ROS/MAP: Religious Orders Study/Memory and Aging Project. ST: spatial transcriptomics.

| Study | Organism | Region | Method | Cell type | significant genes |  |
| --- | --- | --- | --- | --- | --- | --- |
|  |  |  |  |  | R | P-value |
| Habib *et al.*, 2020 | 5xFAD mice * | HP | snRNA-seq | All | 0.26 | 1.10E-09 |
| Miyoshi *et al.*, 2024 | 5xFAD mice * | HP | ST | All | 0.52 | 2.50E-15 |
| Yang *et al.*, 2025 | APP/PS1 mice † | HP | snRNA-seq | CA1 Glut | 0.51 | 7.90E-08 |
|  |  |  |  | CA3 Glut | 0.36 | 0.00044 |
| Gabitto *et al.*, 2024 | Humans (SEA-AD) ‡ | MTG | snRNA-seq | All | 0.093 | 9.90E-07 |
| Mathys *et al.*, 2019 | Humans (ROS/MAP) § | PFC | snRNA-seq | Astro | 0.28 | 0.0067 |
|  |  |  |  | Glut | 0.21 | 2.2E-16 |
|  |  |  |  | GABA | 0.31 | 2.2E-16 |
|  |  |  |  | Oligo/OPC | 0.23 | 2.4E-12 |
|  |  |  |  | Micro/PVN | 0.6 | 0.4 |

\* 5xFAD mice vs WT littermates- References: 4,30
† APP/PS1 mice vs WT - Reference: 31
‡ no AD vs late AD - Reference: 33
§ early vs late pathology - Reference: 32

### Rapid transcriptional changes in glutamatergic and GABAergic neurons of AD-aTNFR1KO

In assessing the transcriptional changes in AD-aTNFR1KO mice with late aTNFR1KO compared to the AD signature, we started with DEG analysis. This analysis identified a substantially smaller number of DEGs distinguishing AD-TNFR1KO from AD mice compared to those distinguishing AD from CTRL mice (**Fig. 3e** and **Supplementary Table 1**). The alluvial plot in **Fig. 3e**, obtained upon aggregating cell populations in pseudobulk, provides a snapshot of the aTNFR1KO-related transcriptional changes and of their relationship with those induced by AD in CTRL mice. The plot highlights three different patterns: (1) “*AD reversal”*, in which a sub-set of the transcriptional changes forming the AD signature is reversed in AD-aTNFR1KO mice. This sub-set comprises prevalently genes that are up-regulated in AD and that undergo down-regulation in the mice carrying the deletion, together with a smaller group of genes that are down-regulated in AD and up-regulated in AD-aTNFR1KO mice; (2) “*AD potentiation”,* which involves a more limited number of genes than those in (1). These genes are either up-regulated or down-regulated in AD and undergo further up- and down-regulation, respectively, in AD-aTNFR1KO mice; the third pattern (3) is “*AD-independent”,* i.e., involves *ex novo* changes in gene expression that are observed in AD-aTNFR1KO mice but are not part of the AD signature. To start deciphering the biological meaning of the transcriptional changes triggered by aTNFR1 deletion in AD mice, and the cell types in which they occur, we resorted again to Reactome pathway enrichment analysis (**Fig. 3f** and **Supplementary Table 3**). In pseudobulk, the pathway most significantly affected by aTNFR1KO was “neuronal system”, followed by pathways more specifically involving synaptic structure and function, membrane excitability, and receptor signaling. When we dissected the signal of these pathways by cell type (Methods), neurons were the almost exclusive source, with the GABAergic neurons family displaying more affected pathways than the glutamatergic family. To verify the specificity of these observations, we used “metabolism”, one of the top pathways targeted by AD (**Fig. 3d**), as negative control and repeated the analysis above. In pseudobulk, we did not find any significant change induced by aTNFR1KO in the 21 metabolism-related pathways; at the level of cell families, we identified few significant changes induced in astrocytic or oligo-OPC’s pathways, but none in neuronal ones (**Extended data, Fig. 5a** and **Supplementary Table 3**). This result supports the specificity of the effect of aTNFR1 deletion in inducing rapid transcriptional changes of synaptic and signaling pathways in glutamatergic and GABAergic neurons. In keeping, a complementary analysis, focusing on cell compartments rather than pathways (Methods, **Fig. 3g** and **Supplementary Table 4**), identified “synapse”, “cell junction” and “plasma membrane” as the top compartments undergoing rapid transcriptional changes upon aTNFR1 deletion in AD mice.

Next, we addressed whether the above transcriptional changes were specific of the AD condition or, instead, deletion of aTNFR1 in CTRL mice could induce similar changes. We thus verified if the top 10 pathways significantly modified in AD-aTNFR1KO mice vs AD mice were also modified in CTRL-aTNFR1KO vs CTRL mice. We found that 4/10 pathways were modified by aTNFR1KO in CTRL mice, but the statistical significance of the effects in CTRL-aTNFR1KO was lower with respect to that in AD-aTNFR1KO mice (**Extended data, Fig. 5b** and **Supplementary Table 3**). We conclude that the transcriptional phenotype involving changes in neuronal synaptic and signaling pathways is, in good part, AD condition-related and cannot be directly reproduced by inducing the aTNFR1KO in CTRL mice.

### Changes in synaptic genes are prominent and different in glutamatergic and GABAergic neurons

We then explored in more detail the results of our pathway analysis aiming to extract an overall biological picture from the transcriptional changes occurring in parallel in glutamatergic and GABAergic neurons, given the required balance in their function for proper memory processing. Therefore, we now considered, for both families, all the pathways significantly modified in AD-aTNFR1KO mice, not just the top 10 ones, as well as the direction of the changes compared to those seen in AD mice vs CTRL (**Fig. 3h** and **Supplementary Tables 2 and 3**). We found that the three general patterns observed in **Fig. 3d** were represented in both neuronal families, but GABAergic neurons showed twice as many modified pathways as glutamatergic neurons (41 vs 20) and a larger quota of AD-independent effects (47% vs 35%), mainly up-regulations. Importantly, 8 of the modified pathways in GABAergic neurons were also modified in glutamatergic neurons, but for 5 of them the modification was in the opposite direction. Among such pathways, “neuronal system”, “neurotransmitter receptors and postsynaptic signal transmission”, and “transmission across chemical synapses” were down-regulated in glutamatergic neurons, and up-regulated in GABAergic ones. Noticeably, the down-regulations in glutamatergic neurons counteracted pre-existing AD-induced up-regulations, whereas the up-regulations in GABAergic neurons occurred *ex novo*. Moreover, selectively in GABAergic neurons, aTNFR1 deletion induced *ex novo* up-regulation of additional pathways related to synapses and receptor signaling, like “protein-protein interaction at synapses”, “neurexins and neuroligins”, “GABA receptor activation”, and “signaling by GPCR”. Overall, the aTNFR1KO induced transcriptional changes appear to target mainly synapse-related processes in both glutamatergic and GABAergic neurons, but with different prominence and often producing opposite regulation.

### Synaptic gene pathways are downregulated in specific glutamatergic neuron subtypes and upregulated in GABAergic ones

To further investigate whether aTNFR1 deletion leads to a coordinated transcriptional “rebalancing” of the AD effects at glutamatergic and GABAergic synapses, we repeated gene set enrichment analysis using SynGO, a framework containing ontology annotation specific to synapse-related genes, locations and processes^34^. Thereby, we identified all the synapse-related DEGs in glutamatergic and GABAergic neurons that are part of the AD signature (AD vs CTRL), or of the aTNFR1 deletion effect in AD mice (AD-aTNFR1KO vs AD). Subsequently, we performed intersectional analysis to understand the relations between these two groups of DEGs (**Fig. 4a** and **Supplementary Tables 5 and 6**). This led us to regroup glutamatergic or GABAergic synaptic DEGs into those belonging to only one groups’ comparison (e.g., up-regulated in AD vs CTRL but not modified in aTNFR1KO vs AD) and those involved in both groups’ comparisons, i.e. the intersectional DEGs (e.g., up-regulated in AD vs CTRL and down-regulated in AD-aTNFR1KO vs AD). Thereby, we confirmed for synaptic DEGs the same three patterns of aTNFR1KO-dependent transcriptional changes identified in the Reactome-based analysis of all DEGs (**Fig. 3d** and **3h**). However, in the case of synaptic DEGs, we found that almost 70% of the expression changes induced by aTNFR1 deletion were *ex novo*, i.e., seen only in AD-aTNFR1KO vs AD. Importantly, they consisted mostly of down-regulated genes in glutamatergic neurons (∼50 DEGs) and up-regulated genes in GABAergic ones (∼80 DEGs). By using an unbiased large language model (LLM)-based contextualized descriptor (Methods), we obtained a functional definition for each of the DEGs’ pools identified by intersectional analysis (**Fig. 4a**). According to such descriptor, the two large pools of down-regulated glutamatergic DEGs and up-regulated GABAergic DEGs, belong to “synaptic signaling” and “synaptic communication”, respectively. We conclude that the opposite transcriptional changes induced by aTNFR1KO deletion at glutamatergic and GABAergic synapses prevalently target molecular actors involved in synaptic transmission or related processes.

**Figure 4.**
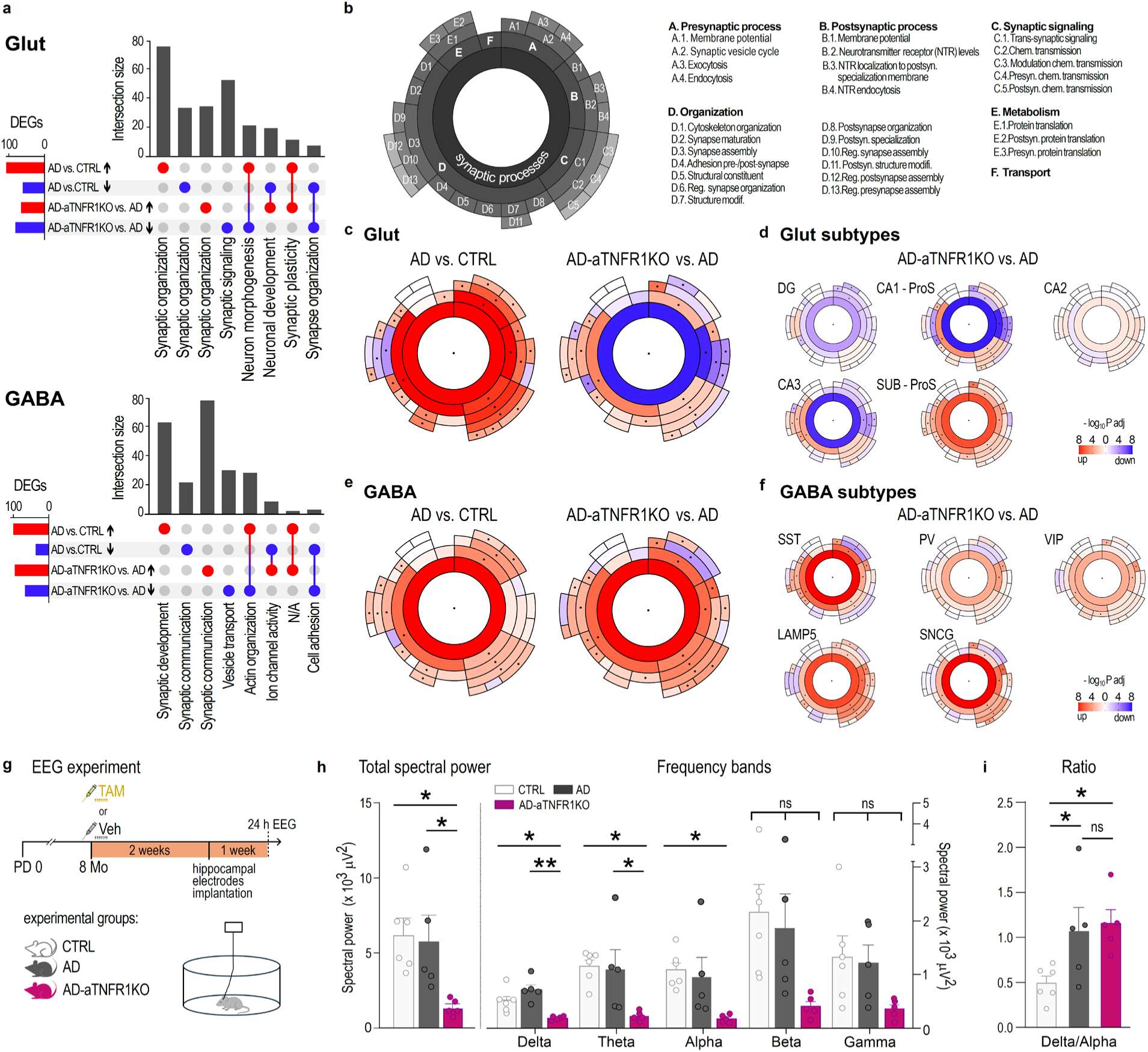
aTNFR1KO downregulates Glut synaptic genes and up-regulates GABA ones, particularly in some Glut and GABA subpopulations, and modifies hippocampal EEG power spectra favoring circuital inhibition. **a**. Upset plots show differentially expressed (blue, down-regulated; red, up-regulated) SynGO-related synaptic genes in AD vs CTRL and AD-aTNFR1KO vs AD group comparisons in glutamatergic (Glut) and GABAergic (GABA) neurons. For each gene set intersection, an LLM-assisted functional summary derived from pathway enrichment analysis is provided below the plots (Methods). **b**. SynGO annotation of sunburst plots. Each “slice” of the plot provides a concentric representation of a synaptic process going from the core process (most internal, dark grey, bold A-F letter) to its sub-hierarchies (progressively more external, lighter grey, letter plus increasing numbers, e.g. A1 to A4). All the ontology terms represented in the sunburst plot and identified by letter plus number are provided in the list adjacent to the plot. **c-d**. Sunburst plot representations of SynGO pathway enrichment analysis for the AD vs CTRL and AD-aTNFR1KO vs AD group comparisons in the Glut neurons family (**c**) and, just for AD-aTNFR1KO vs AD comparison, in each of the hippocampal Glut neuron subtypes (**d**). The heatmap uses an intense red-to-blue gradient scale (in -log_10_Pval) to denote up-regulated (red) and down-regulated (blue) pathways, with black dots in the corresponding sunburst plot section indicating statistical significance. The data show an overall prevailing downregulation of synaptic pathways gene expression in most Glut subtypes in AD-aTNFR1KO vs AD. **e-f**. Sunburst plot representations for group comparisons like in **c-d** but here in the GABA neurons family (**e**) and in each hippocampal GABA neuron subtype (**F**). The data show an overall upregulation of synaptic pathways gene expression in most GABA subtypes. **G**. Schematic representation of the preparation of mice for the EEG experiment. Mouse groups (CTRL, AD, and AD-aTNFR1KO) were treated with Veh or TAM as appropriate (Fig. 1a) at around 8 months of age, and two weeks after the first injection, they were implanted with EEG electrodes in the hippocampus. Following a week of recovery, mice were connected to an EEG system and recorded for 24 h. Analysis was performed on recordings in the 12h-long dark/active phase of the mouse circadian cycle. **h**, *left:* total EEG spectral power analysis shows significantly lower power in AD-aTNFR1KO mice compared to AD and CTRL mice; *right*: analysis in individual frequency bands, shows significantly reduced power in the delta and theta frequencies (total power: one-way ANOVA, p = 0.021; post-hoc tests: AD vs AD-aTNFR1KO: p = 0.040; CTRL vs AD-aTNFR1KO: p = 0.03; CTRL vs AD: p = 0.802; power in each frequency band: one-way ANOVA followed by Holm-Sidak post-hoc tests: delta band: one-way ANOVA: p = 0.014; post-hoc tests: AD vs AD-aTNFR1KO: p = 0.002; CTRL vs AD-aTNFR1KO: p = 0.025; CTRL vs AD: p = 0.078; theta band: one-way ANOVA: p = 0.018; post-hoc tests: AD vs AD-aTNFR1KO: p = 0.038; CTRL vs AD-aTNFR1KO: p = 0.0262; CTRL vs AD: p = 0.078; alpha band: one-way ANOVA: p = 0.0263; post-hoc tests: AD vs AD-aTNFR1KO; p = 0.063; CTRL vs AD-aTNFR1KO: p = 0.0329; CTRL vs AD: p = 0.657; beta band: one-way ANOVA: p = 0.056; gamma band: one-way ANOVA: p = 0.091; for the latter two bands ANOVA was not significant (ns statistical bar), so no post-hoc tests were performed). **i**. In addition to EEG spectral power in each band, the delta/alpha ratio were measured, as higher ratios have been associated with cognitive impairment ^37,38^. The delta/alpha ratio was higher in AD mice compared to CTRL mice but was not reduced in the AD-aTNFR1KO mice (Lognormal one way-ANOVA, p = 0.012; Holm-Sidak post-hoc test: CTRL vs AD: p = 0.037; CTRL vs. AD-aTNFR1KO: p = 0.017; AD vs AD-aTNFR1KO: p = 0.556). N=6 mice for the CTRL group; 5 for the AD group and 5 for the AD-aTNFR1 group, for all analyses.

To have a more complete picture of the glutamatergic and GABAergic synaptic processes and compartments transcriptionally modified in AD (AD vs CTRL) and upon aTNFR1 deletion in AD mice (AD-TNFR1KO vs AD), we performed quantitative pathway enrichment analysis with SynGO (**Fig. 4b-f**). The results are summarized in sunburst charts that provide a snapshot hierarchical visualization of all the annotated synaptic processes and locations (ontology in **Fig. 4b**). In the case of glutamatergic synapses, the sunburst representations in **Fig 4c** show that AD induces widespread up-regulation of both pre- and post-synaptic processes, of synaptic signaling and of structural organization of the synapses. This effect is reversed or largely attenuated by deleting aTNFR1, which induces prevailing downregulation, most notably of post-synaptic processes (all significant changes in **Supplementary Tables 7 and 8**). By applying the same analysis to each individual hippocampal glutamatergic subpopulation identified by snRNA-seq (**Fig. 4d** and **Supplementary Table 8**), the post-synaptic down regulation triggered by aTNFR1 deletion was stronger in the CA1 and CA3 sub-types, less pronounced in the DG subtype, where it was associated with mild down-regulation of pre-synaptic processes and synaptic organization, and was absent in the CA2 and SUB-Pros sub-types, with the latter showing general mild up-regulation. When considering GABAergic synapses (**Fig. 4e** and **Supplementary Tables 7 and 8**), AD prevalently induced transcriptional up-regulation, like at glutamatergic synapses, but on different synaptic aspects, as it predominantly affected presynaptic and structural components. Moreover, aTNFR1 deletion did not reverse these AD-induced changes like in the case of glutamatergic synapses but rather produced new or stronger up-regulations for additional aspects, notably synaptic signaling and post-synaptic processes. In individual GABAergic subpopulations, the above up-regulation was particularly pronounced in the LAMP5 and SNCG sub-types, less in the VIP and PV ones, whereas in SST neurons, pre-synaptic and structural parameters were the most prominently up-regulated (**Fig. 4f** and **Supplementary Table 8**). Overall, SynGO pathway enrichment analysis confirmed that deletion of aTNFR1 triggers a large transcriptional rearrangement at both excitatory and inhibitory hippocampal synapses in AD mice: in specific glutamatergic neuron subtypes, this reverses the widespread synaptic up-regulation produced by AD and produces strong post-synaptic down-regulation; in GABAergic neuron sub-types, instead, aTNFR1 deletion up-regulates notably synaptic signaling and post-synaptic processes. This combined and opposite effect on several molecular determinants of excitatory and inhibitory synaptic transmission suggests that aTNFR1 deletion produces an overall transcriptional upgrading of GABAergic signaling. If translated in synaptic transmission changes such pro-GABAergic resetting could significantly impact on the hippocampal circuit function, which in AD is reportedly shifted towards anomalous hyperexcitation^35,36^.

### AD-aTNFR1KO mice display decreased hippocampal EEG spectral power

To directly verify if and how the transcriptional synaptic resetting triggered by aTNFR1 deletion affects the hippocampal circuit activity in AD mice, we moved to functional studies and recorded EEGs from the hippocampi of 9-months-old CTRL, AD and AD-aTNFR1KO mice, the latter subjected to the late-stage protocol of aTNFR1 deletion (**Fig. 4g**). All mice were recorded for 24h and analysis of total EEG spectral power and power of individual frequency bands was performed during the 12h dark phase/active circadian period (Methods, **Fig. 4h-i**). We found profound differences in the EEGs of AD-aTNFR1KO compared to AD mice, in particular a dramatic reduction in the total spectral power (**Fig. 4h**). This difference was visible also with respect to CTRL mice. At the level of individual frequency bands, the decrease in power with respect to AD was more pronounced and statistically significant for the low frequency bands, particularly for delta. Between AD and CTRL mice we also identified EEG differences, although less prominent: notably, AD mice displayed a combined trend to decreased power for the higher frequency bands and increased power for the delta band, translating into a significantly increased delta/alpha ratio (**Fig. 4i**). This type of EEG alteration has been reported in both AD mouse models^37^ and AD patients^38^, and suggested to indicate impaired cognitive processing. Taken together, the EEG data indicate that aTNFR1 deletion resets the hippocampal circuit toward an inhibition-dominated function, potentially reflecting enhanced GABAergic signaling in line with the transcriptomics data. While this effect may not directly reverse the specific imbalances induced by AD (**Fig. 4i**), the improved memory performance of AD-aTNFR1KO mice indicates that the aTNFR1 deletion-induced circuital changes are beneficial for memory processing.

## Discussion

We here provide the first direct evidence that astrocyte TNFR1 plays a key role in AD pathology and that its deletion, at both pre-symptomatic and symptomatic stages, improves memory performance in aged AD mice. While the beneficial outcome of the early and late interventions is similar, the underlying mechanisms may differ significantly. Thus, when we knocked-out aTNFR1 early in the pathology, the beneficial effect on memory was accompanied by reduced β-amyloid deposits and astrogliosis, suggesting that long-term aTNFR1 ablation may globally attenuate or slow AD pathology and thereby help preserve memory function. However, when we deleted the receptor at late AD stages, the intervention rapidly rescued memory in cognitively impaired 5xFAD mice without, in parallel, reducing β-amyloid deposition or astrogliosis. Therefore, the underlying mechanism in this case must involve changes that can rapidly counteract the causes of the reduced memory performance. Together, the transcriptional resetting of hippocampal synapses and the changed hippocampal EEG spectra, both observed in a time window consistent with the memory rescue, may provide a mechanistic explanation. Thus, the many transcriptional changes triggered by aTNFR1 ablation in both glutamatergic and GABAergic neurons appear to converge mechanistically in targeting synaptic processes, producing simultaneously glutamatergic down-regulation and GABAergic upregulation. This combined action may significantly favour inhibition in the hippocampal circuit, as also indicated by the impressive reduction in EEG power observed in AD mice with aTNFR1KO. Electrophysiological data in both mouse models and humans indicate that subjects with mild cognitive impairment (MCI) or AD often display excitation/inhibition (E/I) imbalances^39^ resulting in hyperexcitation of the hippocampal circuit, including epileptic-like episodes that correlate with impaired cognitive processing^35,36,40–42^. The pro-inhibitory effect of aTNFR1 deletion might thus produce functional rebalancing of the hippocampal circuit excitability, beneficial for memory performance.

Previous work suggests that glial TNFα-TNFR1 signaling exerts an homeostatic control on the hippocampal circuitry via opposing actions at glutamatergic and GABAergic synapses^43^. However, alteration of the TNFα-TNFR1 system, as seen in inflammatory brain conditions, may derange the physiological control and contribute to shifting E/I balance towards excitation^20,44–46^. The fact that aTNFR1 deletion modifies expression of tens-to-hundreds of glutamatergic and GABAergic synaptic genes, suggests that this astrocyte receptor could act as a master regulator of the circuital E/I balance and be key in its derangement in AD. Moreover, the different transcriptional response to aTNFR1 deletion seen in different glutamatergic and GABAergic neuron sub-types hints at a highly sophisticated mode of regulation (and dysregulation in pathology) of hippocampal information processing by aTNFR1.

Several observations in the present study can be relevant to human AD. To start, the transcriptional AD signature across the different hippocampal cell types of our mice correlates with the signature in corresponding cells from human AD patients (**Table 2**), starting from the up-regulation of TNFR1 in astrocytes (**Table 1** and Refs. 4,27). Moreover, the beneficial effect produced by aTNFR1 deletion in our mice is aligned with initial clinical evidence that an anti-TNFR1 agent is of cognitive benefit in a sub-group of AD patients^47^. Memory protection was previously reported in AD mice carrying constitutive TNFR1 knock-out in all cells^48,49^. Thanks to our new transgenic line enabling cell-specific, conditional aTNFR1 deletion, we could critically advance those observations by showing that: (1) memory performance can be improved by deleting TNFR1 solely in astrocytes, a data in line with our previous observations in a mouse model of multiple sclerosis, where we proved that conditional expression of astrocyte TNFR1 in a TNFR1KO background fully reproduced the hippocampal memory phenotype of the pathology^20^; (2) targeting aTNFR1 not only protects memory but also, and crucially, reverses its impairment when triggered at late AD pathology stages; (3) the reversal is fast (assessed by us 3-4 weeks after aTNFR1 deletion) and (4) primarily targets E/I synaptic imbalances, i.e., occurs via a mechanism distinct from those previously considered^48,49^. Together, the above findings shift perspective on TNFα-targeting therapies in AD and other pathologies with cognitive impairment. To start, they indicate that the design of anti-TNFα therapeutics should focus on more specific agents than those tested so far, ideally acting selectively on astrocyte TNFR1. This would avoid long-term inhibition of the receptor in all cells, which likely adds mainly detrimental effects and confuses clinical outcome^7,50^. Secondly, they show that memory performance in late-stage AD mice is not irreversibly impaired and can be rapidly rescued by an *ad hoc* intervention despite the unfavourable environment presenting massive β-amyloid deposits and glial inflammation. This observation is remarkable, as it suggests that - at least until certain stages - memory deficits in AD may largely reflect reversible functional alterations and therefore remain accessible to therapeutic strategies leveraging residual circuital plasticity even in the context of advanced neuropathology. This conclusion is strengthened by the clinical evidence of the existence of “cognitively resilient” AD patients^51^, a minority of subjects who maintain good cognitive capabilities despite presenting massive β-amyloid and tau neuropathology, undistinguishable from that of AD subjects with strong cognitive impairment. Intriguingly, a recent transcriptomics study identifies an abundance of GABAergic neurons, notably of the SST and LAMP5 sub-types^52^, as the most distinctive feature of cognitively resilient subjects, thus indicating that preservation of a robust inhibitory function in memory circuits is key for cognitive resilience. Our findings in AD mice match this observation in human AD, including in identifying the relevance of SST and LAMP5 GABAergic sub-types, which are among those showing the strongest synaptic up-regulation upon aTNFR1 deletion (**Fig. 4f**). Finally, our data identify an unexpected mode of action by which aTNFR1 deletion rescues memory, i.e., rapid functional resetting of the hippocampal circuitry. This mechanism positions aTNFR1-targeting in a class distinct from typical anti-inflammatory interventions^53^, calling for renewed attention to strategies against E/I imbalances. Among them, use of the antiepileptic drug levetiracetam is showing promise in MCI and AD patients, notably in those presenting seizures^54^ Pro-GABAergic approaches are also preconized to be effective^55^ and are under study^56^. Overall, pursuing aTNFR1-targeting may lead to a new class of circuital resetting therapeutics, mechanistically distinct from state-of-art anti-amyloid immuno-therapies^57^, which could thus be used in combination with them to more effectively counteract cognitive decline in AD.

## Extended Data

**Extended Data, Figure 1.**
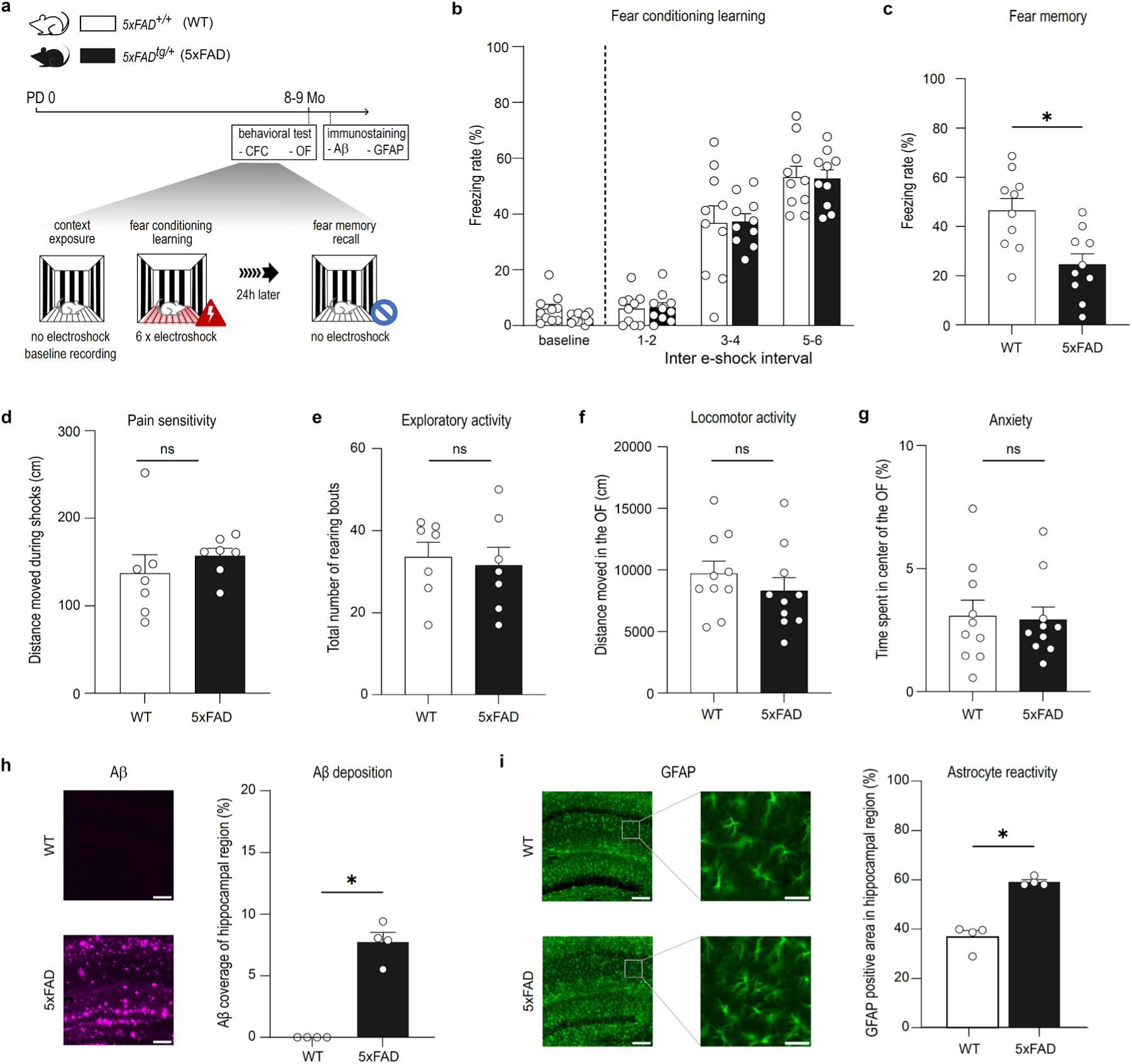
AD pathology phenotyping in native *5xFAD^tg/+^* mice, used for crossing with the *GFAP^CreERT2tg/+^tdTomato^lsl/lsl^TNFR1^fl/fl^* line. **a**. Schematic representation of the AD pathology phenotyping protocol. 8-9 months-old heterozygous mice for the *5xFAD* mutation (*5xFAD^tg/+^,* or 5xFAD for simplicity, black) were compared to littermates lacking the mutation (*5xFAD^+/+^,* or WT - for wild type, white) in the contextual fear conditioning (CFC) and open field (OF) tests. Then, post-mortem, their hippocampi were immunolabeled to reveal β-amyloid deposits and, via GFAP labelling, astrogliosis. **b**. Fear conditioning learning: 5xFAD (black histograms) and WT (white histograms) mice did not differ in their freezing rates during context exposure (baseline) and fear conditioning (Two-way ANOVA, group effect: p = 0.807; time x group interaction: p = 0.800; N=10 mice/group). **c**. Fear memory expression 24 h after conditioning: the 5xFAD group showed significantly lower freezing rates compared to WT mice (Mann-Whitney two-tailed test: U = 17; p = 0.012; N=10 mice/group), confirming the expected memory impairment. **d.** Evaluation of pain sensitivity in 5xFAD vs WT mice, as possible confounder in the freezing performance: the distance moved during the shock period was measured, but the two groups showed no difference (Mann-Whitney two-tailed test: U = 13; p = 0.165; N=6-7 mice/ group). **e**. Likewise, exploratory behavior of 5xFAD vs WT mice was compared based on the number of rearing bouts in each mouse during the context exposure before the first shock (baseline period), showing no group difference (Mann-Whitney two-tailed test: U = 23; p = 0.879; N=7 mice/ group). **f**. Measure of the total distance moved by 5xFAD vs WT mice in the OF test to rule out differences in locomotion affecting freezing rate. The OF test was performed before the CFC test to avoid potential carry-over effects of the test on OF behavior: Again, no differences between the two groups emerged (Mann-Whitney two-tailed test: U = 34; p = 0.248; N=10 mice/group). **g**. Measure of the time spent by the mice in the center of the OF arena, to evaluate anxiety. 5xFAD and WT mice performance was identical (Mann-Whitney two-tailed test: U = 50; p > 0.99; N=10 mice/group). **h**. *left*: representative immunofluorescent images of β-amyloid staining in the hippocampus of WT (*top*) and 5xFAD (*bottom*) mice; Scale bar = 200 µm; *right*: percentage of hippocampal area covered by β-amyloid plaques in the two groups: as expected, staining for β-amyloid (Aβ) was visible only in 5xFAD mice (Mann-Whitney two-tailed test: U = 0, p = 0.029, N=4 mice per group). **i**. *left*: representative immunofluorescent images of GFAP staining in the hippocampus of WT (*top*) and 5xFAD (*bottom*) mice; scale bar = 200 µm or 40 µm for zoomed-in images. *Right:* percentage of hippocampal area covered by GFAP labelling in the two groups: this was significantly higher in 5xFAD mice compared to WT mice, indicating astrogliosis (Mann-Whitney two-tailed test: U = 0; p = 0.029; N=4 mice/group).

**Extended Data, Figure 2.**
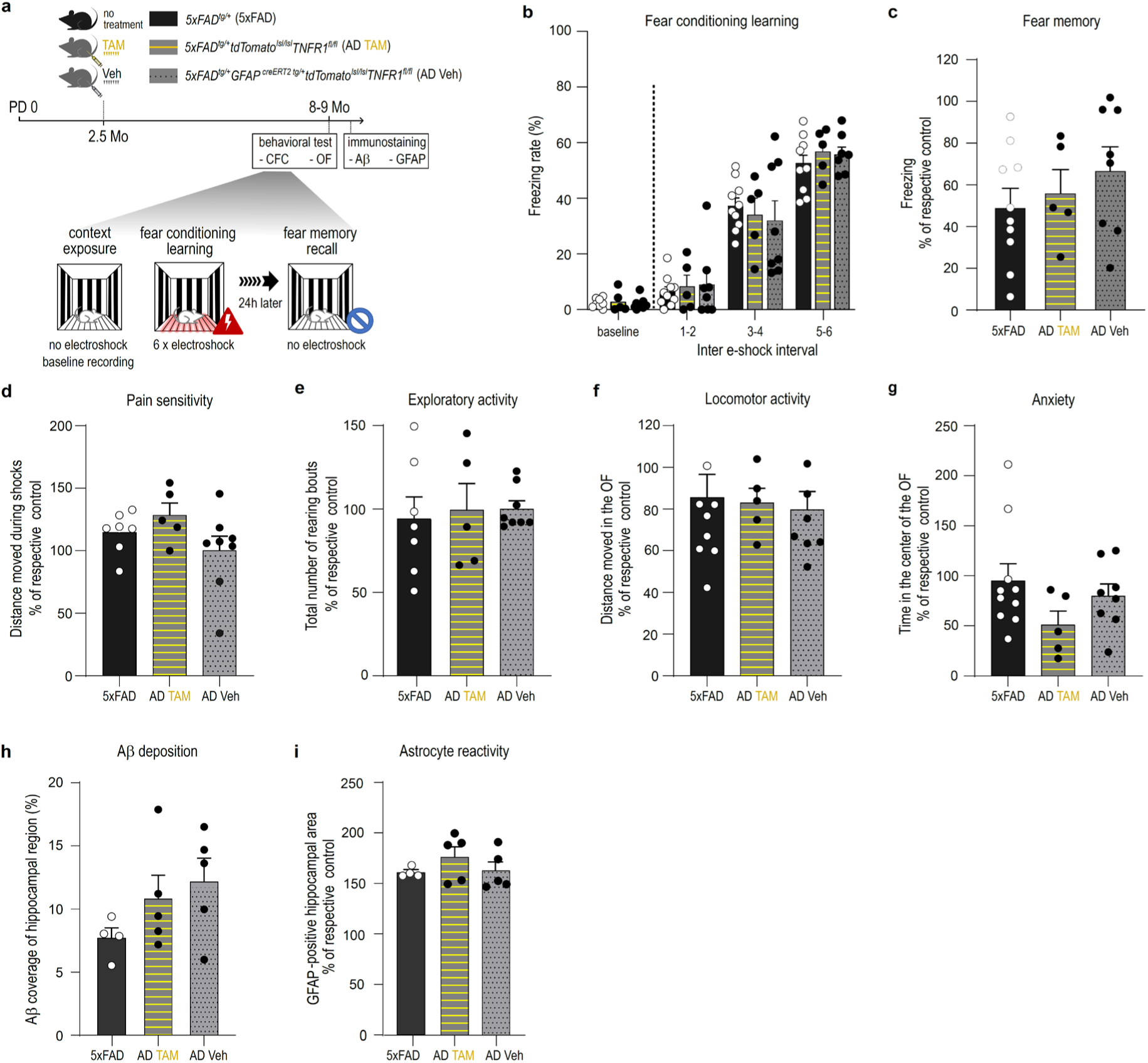
Comparison of AD pathology phenotype shows no difference between native 5xFAD mice and the two derived lines AD Veh and AD TAM. **a**, Schematic representation of the AD pathology phenotyping protocol. At 2.5 months of age, AD Veh (grey, with dotted black line) and AD TAM (grey with yellow line) mice (complete names according to gene inserts given in the scheme and in **Fig. 1a**) were treated with Veh and TAM respectively, whereas native 5xFAD mice (black) were not treated. Mouse groups were compared in the open field (OF) and contextual fear conditioning (CFC) test performance at the age of 8-9 months. Post-mortem, their hippocampi were used for immunohistochemical (IHC) comparative evaluation of Aβ and GFAP immunoreactivity. **b**, Fear conditioning learning: 5xFAD (black histograms), AD Veh (grey with black dots) and AD TAM (grey with yellow stripes) mice did not differ in their freezing rates during context exposure (baseline) and fear conditioning (Two-way ANOVA, group effect, p = 0.989; time x group interaction, p = 0.624; N=5-10 mice/group). **c**, fear memory expression 24 h after conditioning: freezing values for each of the three AD groups (expressing the 5xFAD transgene) are presented as percent of the freezing observed in the respective non-AD littermate control group (not expressing the 5xFAD transgene but treated in the same way) to control for variability between lines and for treatment differences. The percent of control values of the three AD groups are then statistically compared among them. Fear memory relative to respective controls was not different between the three AD groups (Kruskal-Wallis test, H = 1.670; p = 0.434; N=5-10 mice/group). **d,** pain sensitivity measured in AD groups as described in **Extended Data, Fig. 1**, and expressed as percent of control for each group as detailed in **c**, showed no differences between the three AD groups (Kruskal-Wallis test, H = 3.204; p = 0.208; N=5-7 mice/group). **e**, Exploratory behavior, quantified as described in **Extended Data, Fig. 1** and expressed as in **c**, also showed no difference among the AD groups (Kruskal-Wallis test, H = 0.548; p = 0.776; N=5-7 mice/group). **f**, Total distance moved in the OF test relative to respective controls showed no differences between the three AD groups (Kruskal-Wallis test, H = 0.290, p = 0.865, N=5-8 per group). **g**, Time spent in the center of the OF was not different among AD groups, suggesting no differences in anxiety levels (Kruskal-Wallis test, H = 2.801; p = 0.247; N=5-10 mice/group). **h**, percentage of hippocampal area covered by β-amyloid plaques in each of the AD groups, this time expressed in absolute value, was not different among groups (Kruskal-Wallis test, H = 3.586; p = 0.170; N=4-5 mice/group). **i**, percentage of hippocampal area covered by GFAP labeling in each of the AD groups relative to the respective control, was not different among the AD groups (Kruskal-Wallis test, H = 0.966; p = 0.658; N=4-5 mice/group). All comparisons in **c-i** were followed by post-hoc tests between AD Veh and AD TAM groups, without corrections for multiple comparisons in order to allow for a more liberal detection of potential differences between these two groups. Since none of those tests was significant, we pooled the two groups (then called just AD) for all comparisons with CTRL and AD-aTNFR1KO mice.

**Extended Data, Figure 3.**
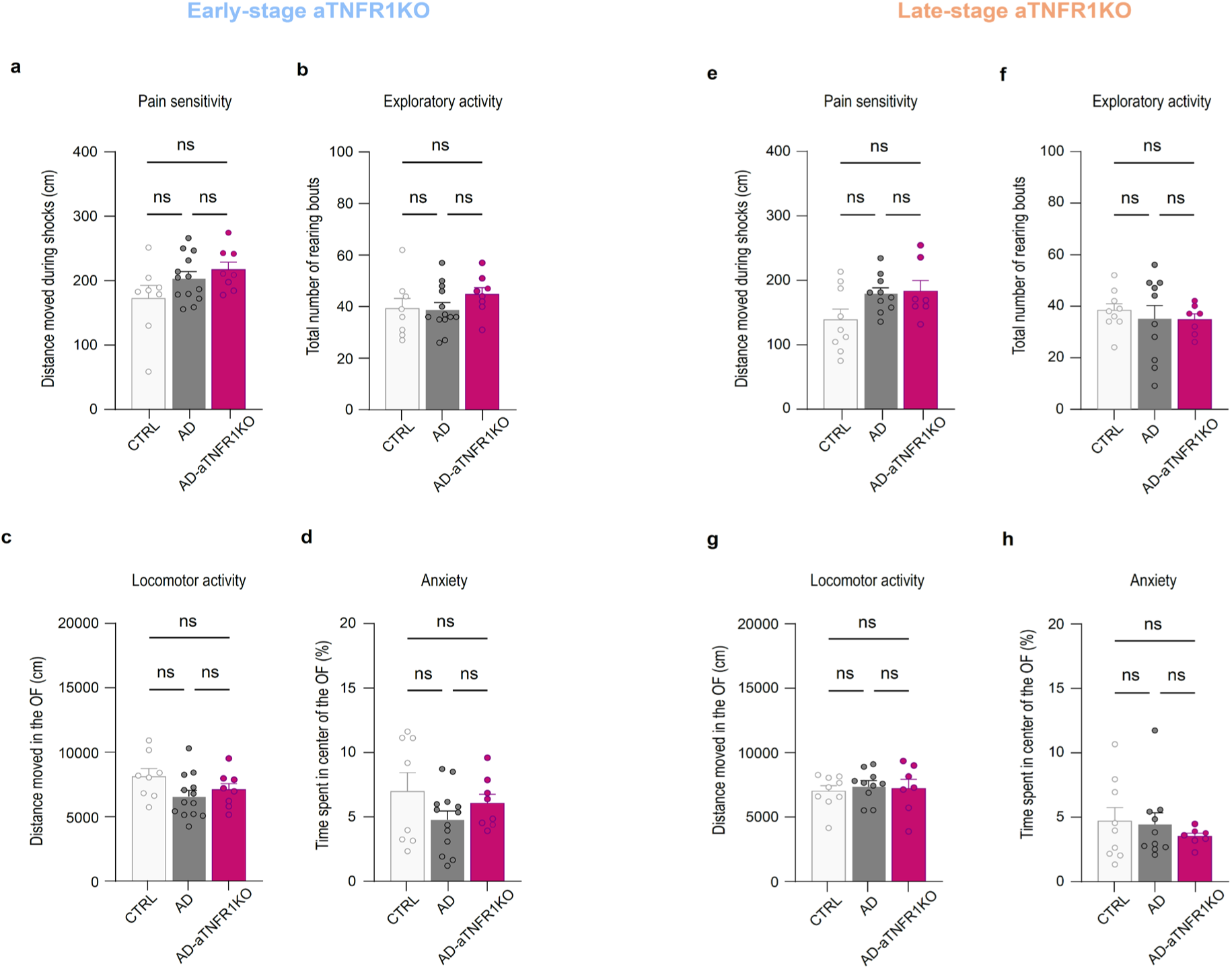
Further behavioural phenotyping of AD-aTNFR1KO mice with either early- or late-stage KO: assessing the contribution of potential confounding factors to the fear memory performance. **a-d**: behavioural phenotyping in 9 months-old mice undergoing the early-stage AD-aTNFR1KO protocol (**Fig. 2a, *left***). The KOs (magenta) are compared to AD (grey) and CTRL (white) mice; tests’ details in Methods and **Extended Data, Fig. 1**; **a,** evaluation of weather a different pain sensitivity to the shocks during CFC contributes to the freezing performance: the results rule out this possibility: pain sensitivity was not different between the three mouse groups (Kruskal-Wallis test: H = 2.818, p = 0.244). **b**, evaluation of differences in exploratory behavior: also for this behavior no difference emerged between the three groups (Kruskal-Wallis test: H = 2.857; p = 0.414). **c,** evaluation of differences in locomotor activity also gave a negative result (Kruskal-Wallis test: H = 3.763; p = 0.288); (**d)** evaluation of anxiety as confounder in the memory performance identified no inter-group difference (Kruskal-Wallis test: H = 3.708; p = 0.295). These results rule out that the above potential confounders contributed to CFC results shown in **Fig. 2b,c**. **e-h**, group comparisons and behavioural phenotyping as in **a-d**, but in 9 months-old mice undergoing the late-stage AD-aTNFR1KO protocol **(Fig. 2a**, *right*). Results were analogous to those in **a-d**, ruling out a contribution to CFC results in **Fig. 2g,h** by the above confounders. None of the measures of: **e,** pain sensitivity (Kruskal-Wallis test: H = 3.871; p = 0.144); **f,** exploratory behavior (Kruskal-Wallis test: H = 0.584; p = 0.747); **g,** locomotor activity (Kruskal-Wallis test: H = 0.323, p = 0.851) and **h**, anxiety (Kruskal-Wallis test, H = 0.117, p = 0.943) revealed intergroup differences. **a-d**, CTRL: N = 8; AD: n = 13; AD-aTNFR1KO: N=8 mice. **e-h**, CTRL: N = 9; AD: N = 10; AD-aTNFR1KO: N=7 mice.

**Extended Data, Figure 4.**
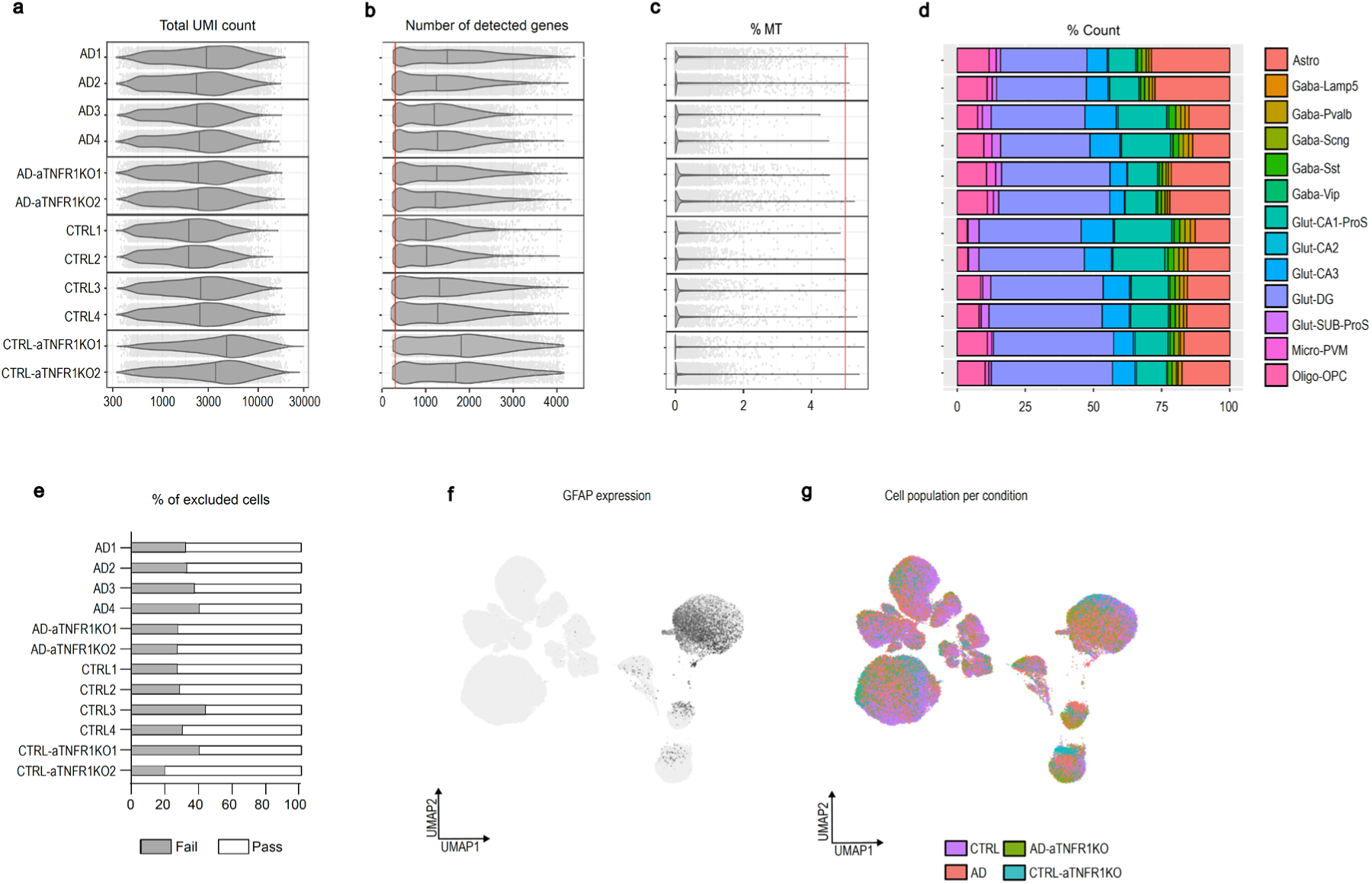
sn-RNA-sequencing quality control. **a**, Total number of unique molecular identifier (UMI) counts per cell in each sn-RNA-seq sample. For differences in the number of samples/group see Methods. **b**, Number of detected genes per cell. Cells in which fewer than 300 genes (red line) were detected were excluded from further analysis. Upper threshold was set at 5000 genes, but no cell reached this level. **c**, Number of mitochondrial (MT) genes per cell. Cells in which more than 5% (red line) of detected UMI were mapped to the mitochondrial genome were excluded from further analysis. **d**, Color-coded comparative visualization of the percentage of the total number of cells (% counts) in each one of our 13 clusters (including astrocytes, oligodendrocytes, microglia and the different glutamatergic and GABAergic subpopulations) for each sample. **e**, percentage of cells that were excluded (grey) or passed (white) quality control and were used for further analysis per sample. **f**, UMAP of GFAP expression showing enriched GFAP expression in astrocytes. **g**, Distribution of cells in each experimental group per cell type cluster. **a-c**, lines in violin plots indicate the median, whiskers the range of data points.

**Extended Data, Figure 5.**
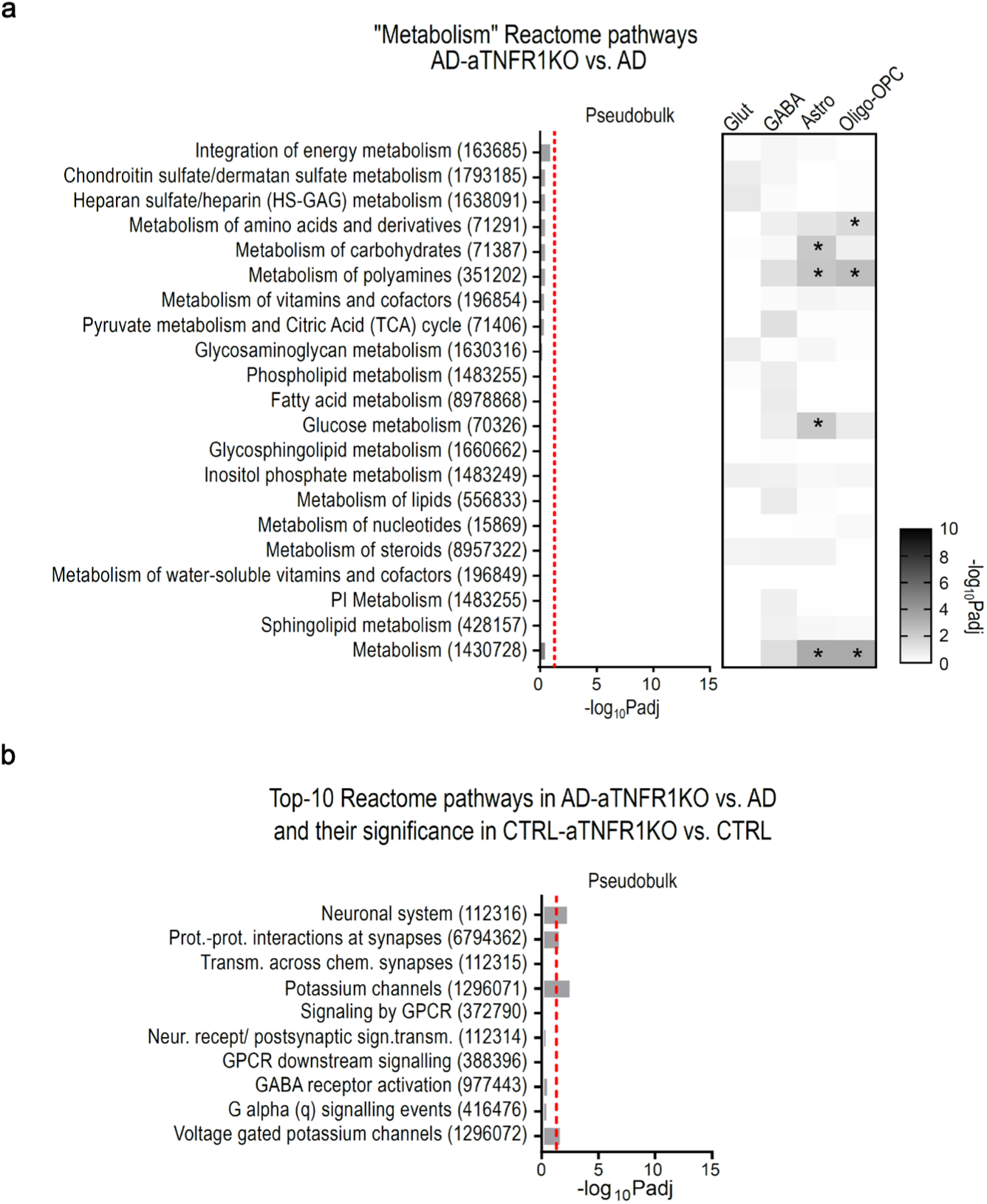
Specificity of the effect of aTNFR1 deletion on the 10 Top Reactome pathways in AD-aTNFR1KO mice and comparison with effect in CTRL-aTNFR1KO mice. **a**, Specificity of the effect of aTNFR1 deletion on Reactome pathways in the AD-aTNFR1KO vs AD groups comparison. To evaluate pathway- and cell type-specificity of the results in **Fig. 3f**, we selected “Metabolism”, one of the Top 10 Reactome pathways differentially regulated by AD vs CTRL (**Fig. 3c**), as negative control, together with its 2^nd^ level subordinated pathways. Pseudobulk analysis found that none of the 21 metabolic pathways was differentially regulated (red dotted line) by aTNFR1KO vs AD. Moreover, the associated heatmap showed significant changes for only few pathways in non-neuronal cells, thus revealing a completely different pattern from the effect identifying the Top-10 regulated Reactome pathways, all in neurons. **b**, AD-specificity of the effect of aTNFR1 deletion on Reactome pathways in the AD-aTNFR1KO vs AD groups comparison. To evaluate if the Top 10 Reactome pathways differentially regulated upon aTNFR1 deletion in AD mice (**Fig. 3f**) were similarly affected when the deletion was produced in CTRL mice, we evaluated their modification in the CTRL-aTNFR1KO vs CTRL group comparison. Pseudobulk analysis revealed that only 4 out of 10 pathways were also significantly regulated (dotted red line) in CTRL-aTNFR1KO mice vs CTRL, albeit with a significance level lower than in AD-aTNFR1KO vs AD mice.

## Methods

### Reagents

The list of reagents utilized in this study is provided as **Supplementary Table 9**.

### Transgenic animals

Mice carrying the tamoxifen-inducible form of Cre recombinase (cre-ERT2) under the human glial fibrillary acidic protein (GFAP) promoter (*GFAPcreERT2; Tg(GFAP-cre/ERT2)1Fki*)^25^ were cross-bred with a conditional tdTomato reporter mouse line for two generations, as previously described^14^. (**Fig. 1a**). The *GFAPcreERT2* gene was maintained in heterozygosis. Subsequently, these mice were crossed with *TNFR1^fl/fl^* mice, provided by George Kollias^26^, to obtain *GFAPcreERT2^tg/+^tdTomato^lsl/lsl^TNFR1^fl/fl^* mice. To produce conditional knockout mice in the context of the AD model, we crossed *GFAPcreERT2t^g/+^tdTomato^lsl/lsl^TNFR1^fl/fl^* mice with *5xFAD APP/PS1* doubly transgenic mice that co-inherit FAD mutant forms of human APP (the Swedish mutation: K670N, M671L; the Florida mutation: I716V; the London mutation: V717I) and PS1 (M146L; L286V) transgenes under transcriptional control of the neuron-specific mouse Thy-1 promoter (line Tg6799 in Jackson Laboratory)^21,62^. Heterozygous *5xFAD^tg/+^* transgenic mice on a C57/BL6-SJL background and age-matched wild-type *5xFAD^+/+^* (WT) controls were used. Since the C57/BL6-SJL strain carries the retinal degeneration Pde6brd1 mutation, which causes visual impairment in homozygosis (Ref. https://www.jax.org/strain/100012), mice were further tested for the presence of the allele. *5xFAD^tg/+^GFAPcreERT2^tg/+^tdTomato^lsl/lsl^TNFR1^fl/fl^* female mice, aged 8-9 months at the beginning of experiments, were used for all experiments. Females were selected for this study because Alzheimer’s disease is more prevalent in women^63^, and female *5xFAD* mice have been reported to have stronger pathology than males^24^. Mice carrying *5xFAD* transgene and their littermate controls (CTRL) were housed together (two to five per cage) in a 12h : 12h light/dark cycle in a temperature and humidity-controlled environment. Food and water access were ad libitum. All experimental protocols in the present study were approved by the Swiss Federal and Cantonal authorities (VD1873, VD2982, VD3876, GE350) and in accordance with the Council directive of the European Union (2010/63/EU). To induce recombination and TNFR1 deletion in astrocytes (AD-aTNFR1KO), *5xFAD^tg/+^GFAPcreERT2^tg/+^tdTomato^lsl/lsl^TNFR1^fl/fl^* mice were intraperitoneally treated once daily for seven days with tamoxifen (TAM, 100 mg/kg, Sigma-Aldrich, Buchs, Switzerland) dissolved in corn oil vehicle (Veh, Sigma-Aldrich) to a concentration of 10 mg/ml. Recombination was induced at either 2.5 months of age, a timepoint when AD cognitive symptoms have not yet appeared (early-stage aTNFR1KO) or at 8 months of age, when cognitive symptoms have been observed (late-stage aTNFR1KO) (**Figs. 2a** and **2g**)^21,22^. Experimental AD mice, i.e., *5xFADt^g/+^GFAPcreERT2^tg/+^tdTomato^lsl/lsl^TNFR1^fl/fl^* treated with appropriate volume of corn oil vehicle (AD Veh) and *5xFAD^tg/+^tdTomato^lsl/lsl^TNFR1^fl/fl^* littermates treated with tamoxifen (AD TAM) were used to test the effect of aTNFR1KO on the AD pathology.

### Genotyping

To detect the floxed or WT TNFR1 (for genotyping) the following primer pair was used: 195 bp (floxed), 134 bp (WT), forward: 5’-CAAGTGCTTGGGGTTCAGGG-3’, reverse: 5’-CGTCCTGGAGAAAGGGAAAG-3’. The PCR conditions for both primer pairs were: 95°C for 3 min, 32 cycles of 95°C for 45 sec, 60°C for 40 sec and 72°C for 1 min 30 sec, followed by an elongation step of 72°C for 10 min. For GFAPcreERT2 genotyping, the following primers were used: 5’-CAGGTTGGAGAGGAGACGCATCA-3’ and 5’-CGTTGCATCGACCGGTAATGCAGGC-3’ with the following conditions: 94°C, 3 min, followed by 35 cycles of : 94 °C, 30 sec,60 °C, 30 sec, 72 °C, 60 sec, and finally 72 °C, 5 min, with the expected product at 500 bp. For the 5xFAD genotyping, 3 primers were used: 5’-CGGGCCTCTTCGCTATTAC-3’, 5’-ACCCCCATGTCAGAGTTCCT-3’, and 5’-TATACAACCTTGGGGGATGG-3’. PCR conditions were: 94°C, 3 min, 10 cycles of (gradient PCR) : 94 °C, 35 sec, 65 °C, decreasing 0.5 °C per cycle, 35 sec, 68 °C, 40 sec, followed by 30 cycles of: 94 °C, 35 sec, 60 °C, 35 sec, 72 °C, 40 sec, followed by 72 °C for 5 min with expected products at 129 bp for the transgenic gene and 216 bp for the wild type. Finally, for the PDE6b gene, the following primers were used: 5’-AAGCTAGCTGCAGTAACGCCATTT-3’, 5’-ACCTGCATGTGAACCCAGTATTCTATC-3’ and 5’-CTACAGCCCCTCTCCAAGGTTTATAG-3’, with conditions: 94°C, 3 min, 10 cycles of (gradient PCR) : 94 °C, 35 sec, 65 °C, decreasing 0.5 °C per cycle, 35 sec, 68 °C, 40 sec, followed by 30 cycles of: 94 °C, 35 sec, 60 °C, 35 sec, 72 °C, 40 sec, followed by 72 °C for 5 min, with expected products at 560 bp for the mutant and 240 bp for the wild type gene.

### Immunofluorescent stainings

At the end of the experiments, the mice were euthanized with an overdose of pentobarbital (150 mg/kg, 1 intraperitoneal (i.p.) injection, Esconarcon, Streuli, Unzach, Switzerland) and transcardially perfused with phosphate-buffered saline buffer (PBS), followed by perfusion with 4% paraformaldehyde (PFA, Electron Microscopy Sciences, Hatfield, PA, USA). Brains were harvested and post-fixated overnight in 4% PFA at 4°C. Afterwards, PFA was removed, and brains were stored in new vials containing a fresh PBS-solution at 4°C. Coronal or sagittal 40 μm-thick sections were cut on a vibratome (Leica, Wetzlar, Germany) and stored in PBS at 4°C until further processing. For evaluation of astrocyte reactive state via GFAP staining and quantification of recombination efficiency (**Fig. 1b, Fig. 2d,f** and **Figure 2i,k, Extended data, Figs. 1-3**), and for staining of β-amyloid deposits (**Fig. 2d-e, Fig. 2i-j, Extended data, Figs. 1-3**), free-floating sections were incubated in a 10% Formic acid solution (Sigma-Aldrich, Buchs, Switzerland) for 10 min at room temperature. Subsequently, sections were washed three times for 10 min in PBS and then incubated for 1 h in blocking solution [foetal bovine serum 10% (Eurobio, les Uils, France), 0.3% triton-x (Merck, Buchs, Switzerland), bovine serum albumin 1mg/ml (Sigma-Aldrich), in PBS]. An overnight incubation at 4°C followed with the following antibodies, all in a concentration of 1:500, in blocking solution: Rat anti-mouse GFAP, (Thermofisher scientific, Geneva, Switzerland, reference number 13-0300) and rabbit anti-pan β-amyloid (Synaptic Systems, Göttingen, Germany, reference number 218103). Sections were then washed three times for 10 min in PBS and incubated for 2.5 h with the following secondary antibodies, in a concentration of 1:1000 in blocking solution: Goat anti-rat Alexa 488 (reference number A11006, Thermofisher), and goat anti-rabbit Alexa 633 (reference number A21071, Thermofisher). Next, sections were washed three times for 10 min in PBS and then incubated in Hoechst 33342 (reference number H21492, Thermofisher, 1:10000) in PBS. Finally mounted with Fluoromount-G mounting medium (Invitrogen, Waltham, MA, USA). Details are provided in **Supplementary Table 9.**

### Imaging and image analysis

For quantification of β-amyloid plaques and GFAP expression, images were acquired on a Zeiss Axio Imager Z1 microscope (Carl Zeiss Microscopy GmbH, Jena, Germany) or an Olympus VS120 slide scanner (Olympus, Basel, Switzerland) using a 10x objective. Laser-excitation wavelength was set at 405 nm for Hoechst 33342, 488 nm with an argon laser for Alexa Fluor 488; and 543 nm and 633 nm with a He/Ne laser for tdTomato and Alexa Fluor 633, respectively. Images were visualized using the ZEN (Blue Edition) software (Carl Zeiss Microscopy GmbH) or VS-ASW (Olympus) and transformed into .tiff format. β-amyloid and GFAP staining were quantified using machine learning-based semantic classification, ilastik 1.4.0^64^. Gaussian smoothing (σ = 3.5), Gaussian gradient magnitude (σ = 1.6), and Hessian of Gaussian eigenvalues (σ = 5.0) were used as feature extractions for β-amyloid positive pixels. Gaussian smoothing (σ = 1.6), Laplacian of Gaussian (σ = 1.0), and Hessian of Gaussian eigenvalues (σ = 3.5) were used as feature extractions for GFAP-positive pixels. All pixels having the prediction of ≥ 60 % to be classified as β-amyloid or GFAP were quantified. To assess recombination in the hippocampal area, cells expressing the reporter gene tdTomato (tdTOM-positive cells) were counted using FIJI (NIH, Bethesda, MD, USA)^65^; the ROI was identified either in the dorsal CA1 stratum radiatum or in the dentate gyrus molecular layer using the freehand selection tool, and Cell Counter plugin in FIJI was used for manual counting. The final cell density is expressed as cells per mm2. In all cases, 1-2 images from 2 slices from 2-7 animals per group were analyzed.

### Preparation of a single-cell suspension from mouse brain regions, FACS isolation of astrocytes and genomic PCR

To verify Cre-dependent recombination, a separate batch of AD, AD-aTNFR1KO and CTRL mice treated with TAM or Veh as appropriate (**Fig. 1a-c, Fig. 2a**), was sacrificed with an overdose of pentobarbital, as mentioned above, and their brains were quickly removed and snap-frozen in isopentane on dry ice and stored at -80 °C until further processing. On the day of preparation, a 1-mm thick coronal section was taken from AD-aTNFR1KO mice, including the cortex and hippocampus, to be used for FACS isolation of astrocytes, and another piece of brain tissue, from individuals from all groups, was taken for whole brain PCR. To FACS isolate astrocytes from frozen brain tissue, a protocol adapted from Ref. ^66^ was used. Briefly, the section was placed in 1 ml of Hibernate-A medium (Thermofisher) and the tissue was thoroughly minced with a razor blade in a glass Petri dish on ice. Afterwards, the solution including the minced tissue was transferred into LoBindTM 1.5 ml Eppendorf tube (Eppendorf, Hamburg, Germany) and centrifuged at 110g for 2 min at 4 °C. The supernatant was discarded and 1 ml of ice cold Accutase solution (Merck) was added and gently mixed. Subsequently, samples were incubated for 30 min at 4°C with head-over-head mixing. Following enzymatic tissue digestion, samples were centrifuged at 960g for 2 min at 4 °C. After removing the supernatant, 0.6 ml of Hibernate-A medium was added and the samples were gently mixed.

Samples were mechanically dissociated through four rounds of gentle trituration using 1.2, 0.8, 0.6 and 0.4 mm syringes. The cell suspensions were then centrifuged at 1700g for 4 min at 4°C. After discarding the supernatant, the pellets were resuspended in Dulbeccòs phosphate-buffered saline (dPBS) containing 1% bovine serum albumin (BSA, Sigma-Aldrich). Finally, samples were filtered through 100-μm pore cell strainers (431752, Corning, Glendale, Az, USA) and the flow through was collected for FACS sorting.

FACS analysis was performed on a BD FACS Chorus system, using a 100 μm nozzle. Compensations were done on single-color control for tdTOM, whereas gates were set on samples from AD mice (not expressing tdTOM). Forward scatter/side scatter gatings were used to remove clumps of cells and debris. For further analysis, 10000 tdTOM-positive and 10000 tdTOM-negative cells were sorted and pelleted with centrifugation at 1700g for 4 min at 4°C. Subsequently, DNA extraction was performed by adding 100 μl of extraction reagent (Quantabio, Beverly, MA, USA), mixed and incubated at 95 °C for 30 min. After the incubation, 100 μl of stabilization buffer were added and samples were stored at -20°C until further processing. DNA samples from whole brain samples without prior FACS sorting were isolated in the same manner.

PCR reactions were performed using the AccuStart II GelTrack PCR SuperMix (Quantbio), as per the manufacturer’s instructions. To identify TNFR1 deletion the following primers were used: TNFR1Δ: 300 bp Forward: 5’-CCTGCAGACACACGGGGAAA-3’, reverse: 5’-TGAACTCAGGTTGCCAGACG-3’. To confirm the presence of the *tdTomato* gene the following primers were used: *tdTomato^lsl/lsl^*: 196 bp, Forward: 5′-GGCATTAAAGCAGCGTATCC-3′; Reverse: 5′-CTGTTCCTGTACGGCATGG-3′. The PCR conditions were: 95°C for 3 min, 35 cycles of 94°C for 30 sec, 61°C for 30 sec and 72°C for 30 sec, followed by an elongation step of 72°C for 5 min. FACS gating strategy and full gel scans are presented in **Supplementary Data**.

### Nuclei isolation

For transcriptomic analyses (**Fig. 3, Fig. 4a-f** and **Extended data, Fig. 4, 5**), mice were sacrificed with an overdose pentobarbital (Esconarcon), the brains harvested and the hippocampi were microdissected and snap frozen on isopentane on dry ice in Eppendorf tubes. Subsequently, they were stored at -80°C until further processing. On the day of nuclei isolation, we pooled 1 hippocampal hemisphere for each 2-3 mice of each experimental group and subsequently resuspended in 2 ml of ice-cold Nuclei EZ buffer (Nuc 101, Merck, Buchs, Switzerland) and gently homogenized by douncing on ice with a Kimble dounce tissue grinder set (D8938, Sigma-Aldrich) (10 times with pestle A and 10 times with pestle B). Following homogenization, the homogenate was transferred to a 5 ml tube, the homogenizer was rinsed with an additional 2 ml of Nuclei EZ buffer, which was transferred to the same 5 ml tube as the homogenate. Samples were gently mixed and placed on ice for 5 min before being centrifuged at 500 g for 5 min at 4 °C. Following centrifugation, the supernatant was removed, the pellet resuspended in 4 ml of ice-cold Nuclei EZ buffer, placed on ice for 5 min and centrifuged again for 500 g for 5 min at 4 °C. Then the supernatant was removed, and the pellet was resuspended in 4 ml of dPBS-1% BSA (Sigma-Aldrich) containing Superase-In RNase inhibitor (Thermofisher) and Enzymatics RNAse inhibitor (Qiagen, Zug, Switzerland), and centrifuged at 500 g for 5 min at 4°C. Finally, the supernatant was removed, the pellet resuspended in 1 ml dPBS-1% BSA containing Superase-In RNase inhibitor (Thermofisher) and Enzymatics RNAse inhibitor (Qiagen) and the nuclei suspension filtered through a 40-μm pore strainer, before proceeding with the following steps.

### TNFR1 expression in published datasets

In order to investigate astrocytic TNFR1 expression in AD (**Table 1**), we searched on-line databases and other published datasets containing gene expression data comparing AD patients to control subjects, or 5xFAD animal models to wild type animals. Most information regarding Refs. 4, 27, and 63-64 was retrieved from https://taca.lerner.ccf.org/ while the information concerning Ref. 59 was retrieved from the supplementary material of the paper, and the information regarding Ref. 58 was retrieved from: http://compbio2.mit.edu/ad_multiregion/.

### Nuclei quantification

20 μl of each sample were stained with Trypan blue and the nuclei were counted on a Nikon AXR system. For nuclei capture and library preparation, 10000 nuclei per sample were loaded on a 10x Chromium X single-cell 3’ kit v.3 RNA sequence platform (10x Genomics, Pleasanton, CA, USA) as per the manufacturer’s instructions. Subsequently, libraries were sequenced on an Illumina NovaSeq 6000 sequencer at a concentration of 1.1 nM. After quantification, each sample was split in two to be used as technical replicates for subsequent steps.

### Pre-processing and quality control steps for sn-RNA-seq data

Sequencing reads obtained in FASTQ format were mapped on the Mus musculus genome primary assembly reference (GRCm39) using 10x Genomics Cell Ranger pipeline (v8.0.0)^67^. Cell Ranger raw matrices outputs were then processed with CellBender (0.3.0)^68^ to remove the signal coming from ambient RNA that is also found in empty droplets. This was done using remove-background command with default arguments to establish the number of epochs needed per sample. A second run of CellBender with the following parameters was then conducted: remove-background with –fpr=0.01, --total-droplets-included=18000, --expected-cells= the “Estimated Number of Cells” from the Cell Ranger summary output file, and the appropriate number of epochs determined from the first iteration. CellBender filtered matrices outputs were then used for downstream analysis in R Statistical Software (v4.3.2). For the analysis, we made an extensive use of Seurat package (v5.0.3)^69^, and Bioconductor packages (release 3.21)^70^ to process the data. More specifically, an initial quality control step consisted in excluding cells with over 5% of mitochondrial reads, or with a number of detected gene per cell outside the range [300-5000, **Extended data, Fig. 4**). A sample was also excluded at this step because it showed QC metrics very different from other samples, in particular the number of cells and distribution of the number of detected genes per cell was highly skewed. In the end, when compared to other samples, this sample had 4 times mores cells not fulfilling our QC criteria. Raw and processed data files generated during the current study have been deposited on the European Molecular Biology Laboratory-European Bioinformatics Institute (EMBL-EBI) repository "ArrayExpress" with accession number E-MTAB-16402.

### Celltype annotation

To annotate cells, a deep neural network (DNN) model was trained on the Allen Brain RNA-Seq Cell Types Database (Mouse Whole Cortex and Hippocampus 10x, https://portal.brain-map.org/atlases-and-data/rnaseq/mouse-whole-cortex-and-hippocampus-10x) to predict cell subclasses from single-cell transcriptomic profile, and was applied to determine identity of our cells. For model training, we used the framework available on https://github.com/BioinfoSupport/scml, that we already used in Ref. 14, and which is optimizing a MultiMarginLoss. Cells annotated with an underrepresented celltype (namely “SMC-Peri”, “VLMC”, “Endo” and “Meis2”), as well as neuronal cell types whose profiles appeared more similar to cortical neurons (as possible contamination from neighboring cortical tissue during dissection) were removed. Also, cells obtaining a low prediction confidence for their celltype were removed (criteria: confidence<1, where confidence is the difference of score between the two top scoring celltypes). Subsequently, doublets were identified using R package scDblFinder^71^ with default parameters.

A preliminary PCA+UMAP dimensionality reduction was then conducted for QC purpose. Using this layout, for each cell, we assessed local consistency of the annotations in PCA space. In detail, for each cell type we calculated the proportion of k-nearest neighbors (k=25) with the same DNN-predicted label and their mean doublet score. Cells with low DNN-predicted consistency (<0.4) or high-doublet score (>0.6) were filtered out. Furthermore, we identified on the UMAP a cluster of cells composed by more than 95% of cells coming from a single sample that we excluded from further analysis.

### Dimensionality reduction

On the remaining quality controlled cells, we ran again Seurat normalization methods (i.e. LogNormalize, FindVariableFeatures, and ScaleData) and performed PCA+UMAP dimensionality reduction. The number of principal components to use was determined by identifying the last PC where the drop in explained variance between consecutive components exceeded 0.1% and retaining PCs up to that point. We then identified clusters and generated visual representation running FindNeighbors, RunPCA, RunUMAP and FindClusters with a resolution of 0.6/1.8. The identified clusters were assigned a cell type by majority vote according to DNN-assigned label of the cells.

### Identification of differentially expressed genes (DEG)

To run differential expression analysis between conditions, cells were aggregated in pseudobulks with method scuttle::aggregateAcrossCells(statistics=”sum”) across samples and cell types, and differential expression analysis was conducted with DESeq2^72^ across conditions (AD vs CTRL, AD-aTNFR1KO vs AD and CTRL vs CTRL-aTNFR1KO) (**Supplementary Table 1**). The differential expression analysis was conducted using two different datasets for the comparison AD vs CTRL; p-values from both datasets were combined with Fisher’s method. This approach was taken because we first performed one run with only two AD and CTRL samples to look into the 5xFAD-dependent transcriptional effects and set up conditions. Subsequently, we repeated the experiment with two samples of each group (also including AD-aTNFR1KO and CTRL-aTNFR1KO groups). A gene was considered significantly differentially expressed when its adjusted-p-value<=0.05. As the number of DEG was imbalanced in certain conditions, due to largely unequal numbers of cells, we considered top 300 upregulated and 300 downregulated genes (ranked by p-value) for pathway enrichment analyses (**Fig. 3f, h, Fig. 4a,c-f**).

For the AD vs CTRL comparison, all significantly differentially expressed genes were used for reactome (**Fig. 3c-d**) and SynGO (**Fig. 4a,c,e**) pathway analyses, since the integration of both experiments allowed much higher statistical power. In the g-profiler analysis for association of different cellular compartments with effects in glutamatergic and GABAergic neurons, the top 300 genes in each direction were merged and used for analysis (**Fig. 3g**). For the sake of clarity, non-canonical or predicted Gm/Rik genes were excluded from all further analyses. In addition, given the low number of microglia cells and the lack of correlation between the gene expression data and previously published datasets (**Table 2**), we excluded this cell type from further analyses.

### Pathway analysis

Pathway enrichment analysis of our DEGs was conducted with databases Reactome^28^ (**Supplementary Table 2 and Supplementary Table 3**) and SynGO^34^ (**Supplementary Table 7 and Supplementary Table 8**), using hypergeometric test implementation available in R package fgsea. Both databases were directly downloaded from their respective website (www.reactome.org, and www.syngoportal.org). For numerical stability and to prioritize general pathways, only pathways with at least 25 annotated genes are tested (method call fgsea::fora(minSize=25)).

The pathways enrichment analyses were performed for each of the 13 identified cell-types, with 5 GABAergic and 5 Glutamatergic subpopulations. To obtain a simplified summary statistic for each of them, we considered for each pathway the smallest enrichment p-value it obtained across the GABAergic (resp. Glutamatergic) subtypes, so that the resulting p-value P obtained could be interpreted as: “pathway X obtained the p-value P in at least one of the GABAergic (resp. Glutamatergic) subclass”. For a better illustration of the variability of the AD transcriptional effects compared to CTRL, the results presented in **Fig. 3d** concerning this comparison only show the uppermost hierarchies and level 1 sub-pathways (reactome database). To summarize the pathways’ results at higher level, we used the same merging strategy of considering the sub-pathway with smallest p-value.

To look into whether the effects seen in glutamatergic and GABAergic neurons in the AD vs AD-TNFR1KO comparison represent a specific effect or a general unspecific transcriptional perturbation we investigated several Reactome pathways related to metabolism as follows: we filtered the column “pathway name” for “metabolism” and column “namespace” selected for “metabolism” and unselected “metabolism of proteins” and “metabolism of RNA”. This selection returned consistently (meaning present in all cell-types) 21 pathways (**Extended Data, Fig. 5** and **Supplementary Table 6**).

In addition, the association between the gene lists in glutamatergic or GABAergic neurons with cellular compartments was done using g:Profiler^73,74^. The full list of top-ranked genes (both up- and down-regulated) per cell type was used as input. The log p-adjusted values were plotted for the top 2 cellular compartments followed by the ones for the major cellular compartments (nucleus, mitochondria, endosome, lysosome, plasma membrane, cytosol, Golgi apparatus, vacuoles and peroxisomes) (**Fig. 3g**). Full analysis results are provided in **Supplementary Table 4**.

### Comparison with previously published datasets

In order to examine whether the transcriptomic differences in our AD vs CTRL comparison were similar to what has been previously described in mouse models of AD and in AD patients (**Table 2**), we compared our data to previously published datasets. First, we accessed GSE143758 (related to Ref. 4), a sn-RNAseq dataset of mouse hippocampus at 7mo across control and 5xFAD experimental groups. For cross-comparison purposes, we re-annotated this dataset using the same neural network used for our data. We then computed a set of gene markers per cell type using Seurat Findmarkers with default parameters, and compared the fold changes obtained with the ones obtained in this study for the comparison AD vs CTRL.

Secondly, we retrieved GSE233208 (related to Ref. 30), a spatial transcriptomic analysis of 5xFAD mouse across ages. From this data, we considered « hippocampus » and « hippocampus-pyramidal » transcriptomic profiles of 8-month animals, and aggregated the profiles to obtain one pseudobulk per sample and conditions. We then looked for DEG between AD and CTRL by running a DESeq2 analysis considering annotations as batches in the design formula. The resulting fold-changes were compared to the average fold-change found in this study between AD and CTRL.

Finally, we considered the list of Human DEG in Ref. ^33^ when comparing cortical scRNA-seq transcriptomic profiles of AD patients against control patients (SEA-AD). FASTQ files containing the sequencing data from the snRNA-seq and snMultiome assays are available through controlled access at Sage Bionetworks (syn26223298). To compare this Human result to our Mouse experiment, we retrieved the list of human-mouse orthologous genes from Ensembl (Release 110), and only considered 1-1 orthologs found between both organisms (n=9407). Then, for each gene, we compared the fold-changes reported in the study with the fold changes found in our dataset between AD and CTRL for the corresponding mouse ortholog. A similar approach was used to compare our dataset with the syn2580853 (related to Ref. 32) and the GSE284797 (related to Ref. 31) datasets. The relevant files for the correlation analysis and associated code, together with the code used in the analysis of our transcriptomic data have been deposited on the Zenodo repository (DOI: 10.5281/zenodo.17911377).

### Large Language Model (LLM)-based functional descriptor analysis

Functional enrichment of gene sets was performed using g:Profiler^73,74^, primarily focusing on the Gene Ontology Biological Process (GO:BP) domain. When fewer than three GO:BP terms were identified for a given gene intersection, the corresponding Molecular Function (GO:MF) terms were also included to ensure sufficient functional representation. All enrichment results were combined across intersections to construct a global term universe, in which each GO term was represented by its minimal adjusted p-value. The global reference was used to provide a biological context for summarizing individual gene intersections. Concise, context-aware functional descriptors were generated using the GPT-4o large language model (OpenAI, 2025) via the official Python SDK (v1.0) implemented in a reproducible conda environment (Python 3.10). For each gene intersection, the model received both the global term universe and the complete list of intersection-specific GO terms and was instructed to output one concise biological phrase (1–3 words), capturing the predominant functional theme. All prompts were executed at a fixed temperature = 0.2 to ensure reproducibility. Token usage, cost, and model metadata were logged, and outputs were compiled into an Excel file (Data S5 and S6). The Python code implementing the LLM-based descriptor workflow is available in Zenodo (DOI: 10.5281/zenodo.17872267).

### Behavioral tests

#### Open Field

To study the effects of astrocytic TNFR1 deletion on locomotion and exploration, 9 months-old native 5xFAD, non-AD littermates (WT), and genetically-modified CTRL, AD and AD-aTNFR1KO mice (see **Fig. 1a**) were tested in the open field (**Extended Data Figs. 1-3**). The open field consisted of a square 45 x 45 cm arena surrounded by 45 cm-high walls. Illumination was set at 30-40 lux. At the beginning of the test, mice were placed in the center of the arena and allowed to explore freely for 20 minutes. Their behavior was monitored and recorded using a video camera mounted on the ceiling above the arena. The total distance moved by the mice during the experiment, their mean speed, and the time that they spent immobile were used as direct measures of assess locomotor activity and exploration using Ethovision XT 11.0 (Noldus, Wageningen, the Netherlands). The time mice spent in a virtual inner zone (12 x 12 cm area in the middle of the arena) was used as inverse index of anxiety. All behavioural apparatuses were carefully washed with 70% ethanol solution between tests.

#### Contextual Fear Conditioning

Mice were subjected to the contextual fear conditioning (CFC) test for associative (US-CS) learning and recall 24 h later^14,75^ (**Fig. 2, Extended Data Figs. 1-3**). The conditioning chamber consisted of a 17 cm x 17 cm arena with contextual cues (black and white stripes) and a grid floor that could be electrified, placed within an isolation chamber (Ugo Basile, Gemonio, Italy), and controlled by Ethovision XT software as above. Mice were first given a 5 min activity test under video recording without electroshocks to assess locomotor activity, baseline freezing and rearing. Next, the fear conditioning learning session started and comprised six inescapable electroshocks (0.2-0.4 mA × 2 sec each), delivered at intervals of 2 min. After the session, the mice were returned to their home cages. The next day (24h later), the mice were placed back into the same arena without electroshocks for recall, i.e. the fear memory recall test. In each of the above sessions, the main measure was the percentage of time spent by the mice freezing during each time interval, with freezing defined as an episode during which no movement of the mouse was detected for at least 2 sec. For the fear conditioning learning session, the mean percentage of time spent by a mouse freezing was calculated for each of the 2-min intervals between the six electroshocks and the cumulative percentage of 2 consecutive inter-shock-intervals (ISI) was summed up and shown (ISI 1–2 min, ISI 3–4 min, ISI 5–6 min). The fear memory recall sessions lasted 11 min, and the mean percentage of time freezing was calculated for minutes 1–10, excluding the first minute of recording. The initial 60 s were omitted because rodents typically display exploratory activity upon re-entry into the context, before recognizing the conditioned environment^76^. Furthermore, the distance that a mouse moved during the electroshock delivery was used as an indicator of pain sensitivity, assuming that such distance is proportional to the sensed pain. All the behavioral apparatus was carefully washed with 70% ethanol solution between tests.

### EEG recordings

#### Stereotactic surgery for electrode implantation

A separate batch of AD, AD-aTNFR1KO and CTRL mice, aged ∼8 months were treated with TAM or Veh once daily for seven days, as mentioned above. Two weeks after the first TAM or Veh injection, surgery for electrode implantation was performed, as previously described^14^. Briefly, mice were anesthetized by isoflurane inhalation in an induction chamber (isoflurane concentration of 2.5 %) and maintained afterwards with an isoflurane concentration of 1-1.5 %. Mice were then mounted on a stereotactic frame (Kopf Instruments, Tujunga, CA, USA). The incision area was shaved and the skin was sterilized with betadine. An injection of 0.2% lidocaine (Streuli, Uznach, Switzerland) (6 mg/kg) and meloxicam (Metacox; Graeub, Bern, Switzerland) (1 mg/kg) was administered locally, 5 min before making a skin incision along the midline. Subsequently, the periosteum was removed to expose the skull. Preemptive buprenorphine analgesia was also administered s.c. (0.1 mg/kg Bupaq, Streuli, Uznach, Switzerland) approximately 30 min before surgery. Six custom-made coated stainless-steel electrodes (AISI316LVM, bare diameter 0.125 mm, coated diameter 0.175 mm, Advent Research Materials, Oxford, UK) were implanted: two between the dura and the skull, bilaterally above the frontal cortex, two intrahippocampally (mediolateral: ±1.8 mm; anterior-posterior: −1.8 mm; dorsoventral: −1.9 mm), and two between the dura and the skull above the cerebellum, one as a reference and one as ground. Electrodes were fixed onto the skull with surgical glue and acrylic dental cement (Paladur, Kulzer, Hanau, Germany) and soldered to an EEG head-mount (Pinnacle Technology, Lawrence, KS, USA). Mice were allowed to recover in their home cage for seven days, while post-operative analgesia was provided (buprenorphine s.c. 0.2 mg/kg twice per day for 2 days and meloxicam if needed).

#### Video and electroencephalography recording analysis

Following post-operative recovery, mice were transferred to a plexiglas cage (Pinnacle Technology) and connected to the EEG recording system, while EEG signal and video were recorded individually from each cage. The EEG signal was continuously recorded at a sampling rate of 250 Hz for 24h (**Fig. 4g**). All recordings contained both light and dark phases under a 12:12 h light cycle (lights on at 7AM; lights off at 7PM). At the end of experiments, mice were euthanized with an overdose pentobarbital i.p. and transcardially perfused with PBS, followed by perfusion with 4% paraformaldehyde (PFA, Electron Microscopy Sciences, Hatfield, PA, USA), and their brains were harvested and stored as mentioned above. Dark-phase (19:00–07:00) intervals were derived from EDF timestamps and analysed as follows: raw EEG signals were imported into Python using MNE-Python and processed in volts to maintain unit consistency. Power spectral density (PSD) for each hippocampal channel was computed using Welch’s method, which estimates frequency content by dividing the signal into overlapping Hann-windowed segments and averaging their periodograms. The resulting PSD values (V²/Hz) were converted to µV²/Hz and integrated across predefined EEG frequency bands—delta (0.5-4 Hz), theta (4-8 Hz), alpha (8-13 Hz), beta (13-30 Hz), and gamma (30-49 Hz)—using Simpson’s rule, yielding absolute band power in µV². Channels with excessive noise, flat traces, or otherwise unphysiological signals were excluded before spectral computation (**Fig. 4h**). For animals with two hippocampal electrodes, PSDs and band-power estimates were calculated separately and averaged to obtain a single representative value per mouse. All analysis code and .edf files are available at Zenodo (DOI: 10.5281/zenodo.17868880).

### Statistics and reproducibility

Graphpad prism (versions 8 or 9, Dotmatics, Boston, MA, USA) was used for all statistical analyses, except for bioinformatic analyses for transcriptomics for which R (R Core Team, 2023) was used. Data are presented as mean ± s.e.m., unless stated differently in legends, and normalized data are calculated relative to baseline, unless otherwise indicated. Data were considered to be significantly different when P < 0.05. P values less than 0.05, 0.01, and 0.001 are indicated in figures by 1, 2, and 3 asterisks, respectively. For comparison of gene expression between our dataset and previously published datasets, Pearson’s correlation was used. The two-tailed Mann-Whitney test was used when comparing the means of two independent experimental populations. The Kolmogorov–Smirnov test was used when comparing cumulative distributions between two independent experimental populations. ANOVA or Kruskal-Wallis tests were used when comparing the means of more than two populations. Two-way repeated measures ANOVA with or without independent treatment groups was performed on time-course experiments. For cases in which ANOVA or Kruskal–Wallis tests yielded significant effects, appropriate post hoc comparisons were used to identify significant pairwise differences: Dunn’s multiple comparisons test following Kruskal–Wallis, the Holm–Šídák test after one-way ANOVA, and either Tukey’s multiple comparisons test (for three groups) or the Šídák multiple comparisons test (for two groups) after two-way ANOVA. When liberal exploratory pairwise comparisons were required despite non-significant omnibus effects, uncorrected Fisher’s LSD or uncorrected Dunn’s tests were applied. For the comparison of Delta/Alpha band ratio among groups, the lognormal one-way ANOVA test was used, followed by Holm–Šídák post-hoc tests. Grubbs outlier test was performed which led to the exclusion of statistical outliers in some cases. For transcriptomic analyses, pseudobulk count data were modelled using DESeq2^72^, and differential expression was assessed using a negative binomial generalized linear model with Wald statistics. Pathway enrichment was evaluated using a hypergeometric overrepresentation test, and p-value adjustment for multiple-testing was performed using the Benjamini–Hochberg FDR method. In Pathway enrichment panels in figures, the -log_10_Padjusted is plotted. Heatmaps indicate -log_10_Padjusted levels and asterisks or dots denote statistical significance in each cell type. Data shown from representative experiments were repeated with similar results in at least two independent biological replicates, unless otherwise noted. Wherever possible, individual datapoints were plotted and exact sample sizes are mentioned in the legend. Sample sizes were estimated empirically based on previous studies. Full output of each statistical analysis is included in **Supplementary Table 10**.

## Data availability statement

The new single-nucleus RNA-seq datasets generated during the current study are available in the European Molecular Biology Laboratory-European Bioinformatics Institute (EMBL-EBI) repository "ArrayExpress" with accession number E-MTAB-16402. The already available datasets analysed during the current study and their public domain resources are indicated in the reporting summary form and in Methods. Associated files have been deposited in Zenodo (10.5281/zenodo.17911377). EDF files containing the EEG recordings generated in the present study have been deposited in Zenodo (10.5281/zenodo.17868880). The other datasets are available from the corresponding authors on reasonable request.

## Code availability statement

Algorithm and codes used to analyse single-nucleus RNA-seq datasets are described in Methods and Reporting Summary form and are available in Zenodo repository (10.5281/zenodo.17911377), together with the code used for the correlation analysis presented in **Table 2**. The folder "code" is an archive file containing an R project including all scripts to reproduce the snRNA-seq analysis. The folders “habib”, “miyoshi”, “gabitto” and “mathys” contain custom R code used for gene expression correlation analyses **(Table 2**). For each external study, the corresponding output files are provided as separate compressed folders, including TSV files summarizing correlation coefficients used to generate the figures in the manuscript. The R Markdown files contain extensive comments describing the purpose of each function, data transformation, and statistical test. The Python code used for the unbiased LLM analysis presented in **Figure 4a** is available in Zenodo (10.5281/zenodo.17872267) together with a README file describing the analysis workflow and required dependencies. The Python code used for EEG data processing and analysis has been deposited in Zenodo (10.5281/zenodo.17868880). Documentation for the data and code is provided in separate README files in the same repository. Codes used to analyse β-amyloid and GFAP immunoreactivity areas are described in Methods and public domain resources are listed in **Supplementary Table 9.**

## Acknowledgements

Research in the Volterra lab was supported by Swiss National Science Foundation (SNSF) Grant/Award Number 31003B-201276; Stiftung Synapsis - Demenz Forschung Schweiz DFS, Grant/Award Number: 2018-PI-01 and by Wyss Center for Bio and Neuroengineering, Geneva. The authors thank George Kollias and Frank Kirchhoff for providing the *Tnfr1^fl/fl^* and *GFAP^CreERT2^*mouse lines, respectively; Tara Canonica for IHC experiments, imaging and quantification in the initial part of the project; Elisabeth Urban and Theo Ribierre for FACS isolation and nuclei counting at the NeuroNA Human Cellular Neuroscience platform (Campus Biotech, Geneva), Sergi Roig Puiggros for consultation regarding nuclei isolation; The iGE3 Genomics platform of the University of Geneva for single-nucleus RNA-seq sequencing; Cheng-Hsi Wu and Léa Camelot for preliminary patch clamp experiments at hippocampal synapses; Christopher Pryce and Fulvio Magara for advice on protocols and analysis of behavioural experiments; Denis Jabaudon and Ron Stoop for critical reading of the manuscript.

## Author contributions

**TK** participated to the design of the project, supervised most of its experimental and analytical parts, designed crossings and supervised breeding schemes and maintenance of the transgenic lines with **RdC** and **IZ**, designed, performed and analyzed morphological and behavioral experiments, participated in single-nucleus transcriptomics and EEG experimental design and analysis, including writing in-house codes for data analysis, wrote parts of the manuscript, designed most figures, and supervised some co-authors during analyses and writing of the paper; **IZ** designed, performed and analyzed EEG experiments, participated in single-nucleus transcriptomics experimental design, performance and analysis, performed and analyzed morphological experiments, supervised maintenance of the transgenic lines with **TK**, prepared cell suspensions for FACS-isolation, performed genomic PCR experiments, prepared related figures and wrote related manuscript’ sections and contributed to the overall manuscript writing; **JP** designed and performed bio-informatic analysis of snRNA-seq experiments, in part supervising **AA**, prepared the related figures and wrote the corresponding ms. sections; **AA** performed bio-informatic analysis of existing databases and participated in analysis of sn-RNA-seq experiments; **RdC** designed crossings and supervised breeding schemes and maintenance of the transgenic lines with **TK** and **IZ** and participated in IHC and behavioural experiments in the initial part of the project. **MM** participated in the design and performance of electrophysiological experiments, in the writing of the manuscript, and gave strategic input on the project. **LT** advised on the strategy, experimental performance and analysis of the transcriptomics part of the project, including figures design and writing of the ms text. **AV** supervised the entire project, defined its strategy and, together with **TK** and other co-authors, designed its different components and wrote the manuscript.

## Additional information

Supplementary Information is available for this paper in the form of 10 Supplementary Tables,1 Supplementary Data File and 1 Source Data Table.

**Supplementary Table 1**: list of differentially expressed genes in pseudobulk and individual cell types/families across comparisons presented in **Fig. 3c** and **3e**. Log_2_-transformed fold changes are provided. Differential expression was assessed using a negative binomial generalized linear model with Wald statistics. P-values and adjusted P-values (false discovery rate) are also provided.

**Supplementary Table 2**: results of the Reactome pathway enrichment analysis based on two independent experiments for the AD vs CTRL groups comparison in pseudobulk and individual cell types/families (**Fig. 3d)**. Pathway enrichment was evaluated using a hypergeometric overrepresentation test, and p-value adjustment for multiple-testing was performed using the Benjamini–Hochberg FDR method.

**Supplementary Table 3**: results of the Reactome pathway enrichment analysis for the AD-aTNFR1KO vs AD and CTRL-KO vs CTRL group comparisons in pseudobulk and in all cell types/families presented in **Figs. 3f, 3h** and **Extended Data Fig. 5a,b**. Pathway enrichment was evaluated using a hypergeometric overrepresentation test, and p-value adjustment for multiple-testing was performed using the Benjamini–Hochberg FDR method

**Supplementary Table 4**: results of g:Profiler analysis for cellular compartment enrichment presented in **Fig. 3g**. Separate lists of genes and of results of the g:Profiler analysis are provided for glutamatergic and GABAergic neuron families.

**Supplementary Table 5**: SynGO-related lists of differentially expressed genes for glutamatergic and GABAergic neuron families used for the functional pathway enrichment analysis presented in **Fig. 4a**. Each list corresponds to a specific gene intersection and includes only genes annotated in the SynGO database. Gene identifiers are provided as official gene symbols.

**Supplementary Table 6:** results of SynGO Gene Ontology/Biological processes enrichment (GO:BP) analysis for each gene intersection presented in **Supplementary Table 5**. For each intersection, enriched GO terms and their corresponding adjusted p-values (false discovery rate corrected) are reported. These tables served as direct input for the LLM-based functional descriptor analysis presented **Fig. 4a**.

**Supplementary Table 7:** results of SynGO pathway enrichment analysis for the AD vs CTRL groups comparison in glutamatergic and GABAergic neuron families presented in **Figs. 4c** and **4e**. Evaluation was done using a hypergeometric overrepresentation test, and p-value adjustment for multiple-testing was performed using the Benjamini–Hochberg FDR method.

**Supplementary Table 8**: results of SynGO pathway enrichment analysis for the AD-aTNFR1KO vs AD groups comparison in glutamatergic and GABAergic neuron families and in specific glutamatergic and GABAergic sub-types presented in **Figs. 4c-f**. Evaluation was done using a hypergeometric overrepresentation test, and p-value adjustment for multiple-testing was performed using the Benjamini–Hochberg FDR method.

**Supplementary Table 9**: List of all the reagents and tools that were used in this study.

**Supplementary Table 10**: Statistical details for all the analyses presented in the Figures and Extended Data Figures of our study.

**Supplementary Data:** Full gel images and FACS gating

**Source Data Table:** Source data presented in all the Figures and Extended Data Figures.

**Correspondence and requests for materials** should be addressed to Andrea Volterra and Toko Kikuchi.

